# Caspases switch off m^6^A RNA modification pathway to reactivate a ubiquitous human tumor virus

**DOI:** 10.1101/2020.11.12.377127

**Authors:** Kun Zhang, Yucheng Zhang, Yunash Maharjan, Febri G Sugiokto, Jun Wan, Renfeng Li

## Abstract

The methylation of RNA at the N6 position of adenosine (m^6^A) orchestrates multiple biological processes to control development, differentiation, and cell cycle, as well as various aspects of the virus life cycle. How the m^6^A RNA modification pathway is regulated to finely tune these processes remains poorly understood. Here, we discovered the m^6^A reader YTHDF2 as a caspase substrate via proteome-wide prediction, followed by *in vitro* and *in vivo* validations. We further demonstrated that cleavage-resistant YTHDF2 blocks, while cleavage-mimicking YTHDF2 fragments promote, the replication of a common human oncogenic virus, Epstein-Barr virus (EBV). Intriguingly, our study revealed a feedback regulation between YTHDF2 and caspase-8 via m^6^A modification of *CASP8* mRNA and YTHDF2 cleavage during EBV replication. Further, we discovered that caspases cleave multiple components within the m^6^A RNA modification pathway to benefit EBV replication. Together, our study establishes that caspase disarming of the m^6^A RNA modification machinery fosters EBV reactivation.

Teaser

Cellular m^6^A RNA modification machinery is cleaved by caspases to foster the reproduction of a common human tumor virus

## Introduction

Epstein-Barr virus (EBV) is a ubiquitous tumor virus causing several types of cancer of B cell and epithelial cell origin (1, 2). Globally, EBV infection causes more than 200,000 new cancer cases and 140,000 deaths per year (3). The life cycle of EBV include a quiescent latent phase and an active replication phase (4). The switch from latency to lytic replication, also called reactivation, involves a series of signaling pathways that drive the expression of two EBV immediate early genes, *ZTA* and *RTA* (5). Host factors that restrict or promote the expression of *ZTA* and *RTA* determine the threshold for EBV lytic cycle activation (5–8). Oncolytic therapies based on reactivation of latent virus is a promising approach for targeted treatment of EBV-associated cancers (9), but these strategies require a deeper understanding of how the EBV life cycle is dynamically regulated by key cellular processes and pathways.

The N6-methyladenosine (m^6^A) modification of viral and cellular mRNAs provides a novel mechanism of post-transcriptional control of gene expression (10–12). M^6^A modification is dynamically regulated by methyltransferases (writers, METTL3/METTL14/WTAP/VIRMA) and demethylases (erasers, ALKBH5 and FTO) (10, 13–15). The m^6^A-specific binding proteins (readers, e.g. YTHDF1/2/3 and YTHDC1/2) regulate various aspects of RNA function, including stability, splicing and translation (10, 11, 16, 17). YTHDF2 mainly regulates mRNA decay through direct recruitment of the CCR4-NOT deadenylase complex (16, 18). Recent studies showed that YTHDF1 and YTHDF3 share redundant roles with YTHDF2 in RNA decay (19, 20). All three YTHDF proteins undergo phase separation through binding to m^6^A modified RNA (21–23).

The m^6^A RNA modification pathway has been shown to promote or restrict herpesvirus infection and replication depending on the viral and cellular context. The depletion of YTHDF1 and YTHDF2 has been shown to promote EBV lytic protein expression (24, 25). EBV immediate-early protein ZTA suppresses METTL3 expression through binding to its the promoter (26) and EBV latent protein EBNA3C activates METTL14 transcription and directly interacts with and stabilizes METTL14 to promote oncogenesis (25). Both EBV latent and lytic genes are modified by m^6^A during primary infection, latency or lytic reactivation (25, 27). In one study, it was shown that YTHDF2 binding to m^6^A-modified viral RNA restricts Kaposi’s sarcoma-associated herpesvirus (KSHV) and EBV replication through promoting RNA decay (25, 28). However, another study showed that YTHDF2 either restricts or promotes KSHV replication depending on cell types (29). In addition, YTHDC1 was shown to bind to m^6^A-modified KSHV *RTA/ORF50* to regulate its pre-mRNA splicing and promote lytic replication (30). SND1 was recently discovered as another m^6^A reader that binds and stabilizes KSHV *RTA/ORF50* transcript to promote KSHV replication (31). M^6^A modification of cellular genes promotes human cytomegalovirus (HCMV) replication through downregulating the interferon pathway (32). The complex function of m^6^A pathway in herpesvirus infection suggests that this pathway must be finely regulated during viral infection or reactivation in different cellular context.

Although caspase-mediated cell death is intrinsically hostile for viral replication, emerging studies from our group and others have demonstrated that caspase cleavage of cellular restriction factors promote EBV and KSHV reactivation (6, 7, 33–35). Based on two evolutionarily conserved cleavage motifs we discovered for PIAS1 during EBV reactivation (7), we screened the entire human proteome and identified 16 potential caspase substates that carry the same motifs. Among these proteins, we validated 5 out of 6 as *bona fide* caspase substrates and then focused on the m^6^A reader YTHDF2 and, subsequently, the entire m^6^A RNA modification pathway in the context of EBV reactivation process. We found that caspase-mediated cleavage converts YTHDF2 from a restriction factor to several fragments that promote EBV reactivation. Mechanistically, we demonstrated that YTHDF2 promotes the degradation of *CASP8* mRNA via m^6^A modification to limit caspase activation and viral replication. Importantly, we further illustrated that multiple m^6^A pathway components, including methyltransferases and other readers, were also cleaved by caspases to promote viral replication. Together, these findings uncovered an unexpected cross-regulation between caspases and the m^6^A RNA modification pathway in switching EBV from latency to rapid replication.

## Results

### Cleavage sequence-guided screening identifies potential novel host factors involved in EBV replication

Recently, our group demonstrated that PIAS1 is an EBV restriction factor that is cleaved by caspase-3, -6 and -8 to promote EBV replication (7). The two evolutionally conserved sequences surrounding PIAS1 cleavage site D100 and D433, (LTYD*G and NGVD*G), are quite different from the canonical caspase cleavage motifs (36). We predicted that other proteins within the cellular proteome may contain the same sequences and hence are potentially regulated by caspases. Using these two cleavage sequences to search the human proteome, we discovered 16 additional putative caspase substrates, including RNA binding proteins (YTHDF2 and EIF4H), chromatin remodeling proteins (MTA1 and EHMT2) and a SARS-CoV2 receptor ACE2 (**Figure 1A** **& 1B**).

**Figure 1.**
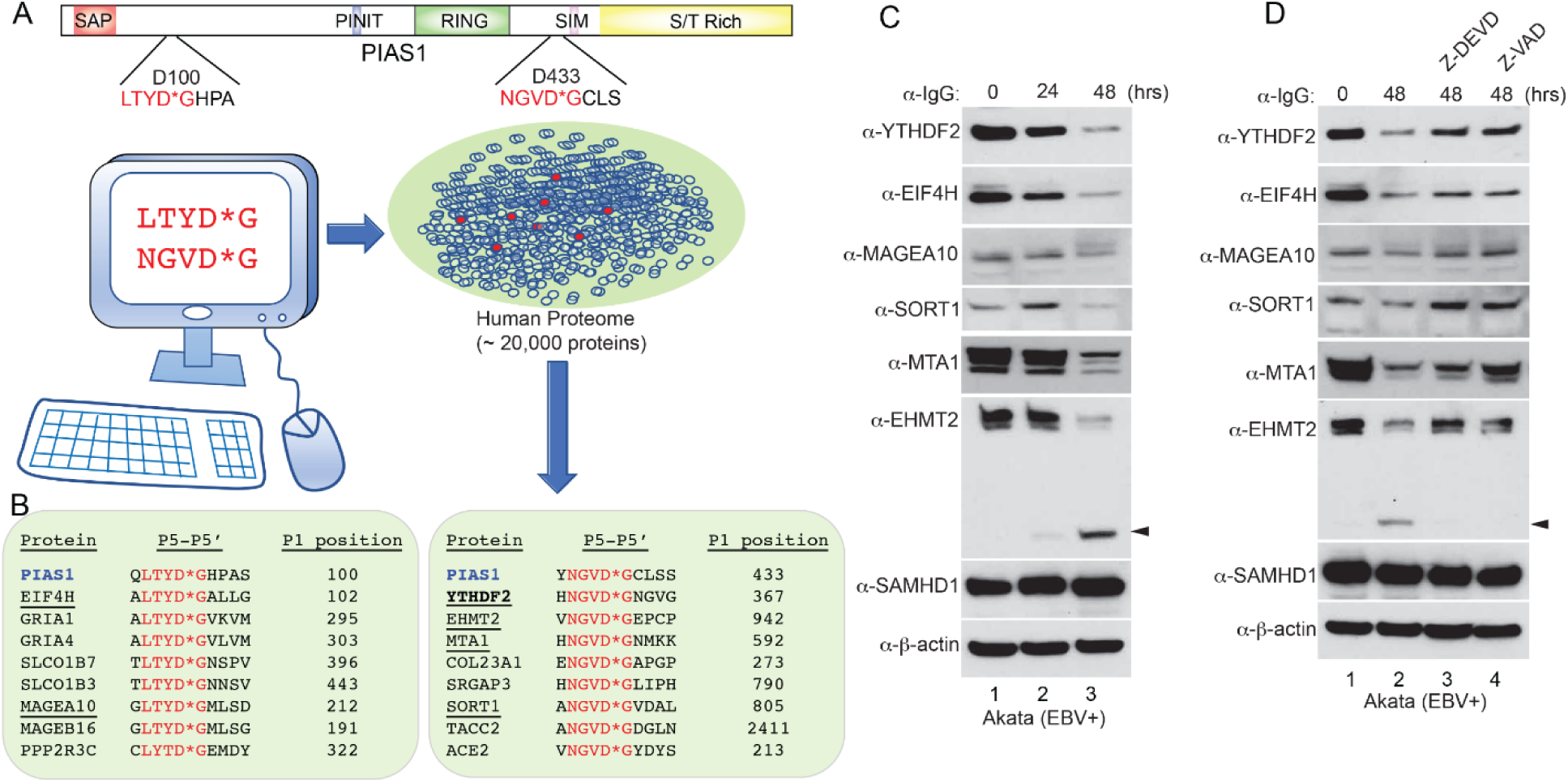
Virtual screen identifies new potential caspase substrates during EBV reactivation. (A) The cleavage motifs derived from PIAS1 (LTYD*G and NGVD*G) were used to virtually screen the entire human proteome for proteins sharing the same sequences. The human proteome dataset containing approximately 20,000 human protein-coding genes represented by the canonical protein sequence was downloaded from UniProtKB/Swiss-Prot. (B) 16 additional proteins were extracted from the screen. 8 proteins carry the LTYD*G motif (left) and 8 proteins carry the NGVD*G motif (right). 6 proteins (underlined) were selected for further validation. (C) Protein downregulation during EBV reactivation. Akata (EBV+) cells was treated with anti-IgG antibody to induce EBV reactivation for 0, 24 and 48 hrs. Western Blot showing the downregulation of 6 selected proteins using antibodies as indicated. SAMHD1 and β-actin were included as controls. Arrowhead denotes the cleaved fragment for EHMT2. (D) Caspase inhibition blocks the degradation of YTHDF2, MAGEA10, SORT1 MTA1 and EHMT2. The Akata (EBV+) cells were either untreated or pretreated with a caspase-3/-7 inhibitor (Z-DEVD-FMK, 50 μM) or pan-caspase inhibitor (Z-VAD-FMK, 50 μM) for 1 hr, and then anti-IgG antibody was added for 48 hrs. Western Blot showing the protein levels of 6 selected proteins using antibodies as indicated. SAMHD1 and β-actin were included as controls. Arrowhead denotes cleaved EHMT2 fragment.

We selected 6 candidates (YTHDF2, EIF4H, MAGEA10, SORT1, MTA1, and EHMT2) to monitor their stability in EBV-positive Akata cells upon anti-IgG-induced B cell receptor (BCR) activation, a process that triggers caspase activation and EBV reactivation (**Figure 1C**) (6, 7, 37). All 6 proteins were downregulated upon BCR activation but not SAMHD1, a cellular restriction factor against EBV replication (38). To further determine whether any of these proteins are regulated by caspase, we pre-treated the cells with a caspase-3/7 inhibitor or a pan-caspase inhibitor followed by IgG-crosslinking of BCR. We found that the downregulation of YTHDF2, MAGEA10, SORT1, MTA1, and EHMT2 was reversed by the caspase inhibition, suggesting that these proteins are *bona fide* caspase substrates in cells (**Figure 1D**). Caspase inhibition partially restored the EIF4H, indicating an involvement of other pathways in controlling EIF4H protein level (**Figure 1D**).

To determine whether any of these proteins play a role in EBV life cycle, we utilized a CRISPR/Cas9 genome editing approach to knock out the corresponding genes in Akata (EBV+) cells, and then evaluated the viral lytic gene products upon BCR stimulation. The depletion of YTHDF2 and EIF4H promoted EBV ZTA and RTA protein expression (**Figures 2A, 2B & S1A**). In contrast, the depletion of MAGEA10, SORT1, EHMT2 and MTA1 did not affect EBV protein expression (**Figures S1B-S1E**).

**Figure 2.**
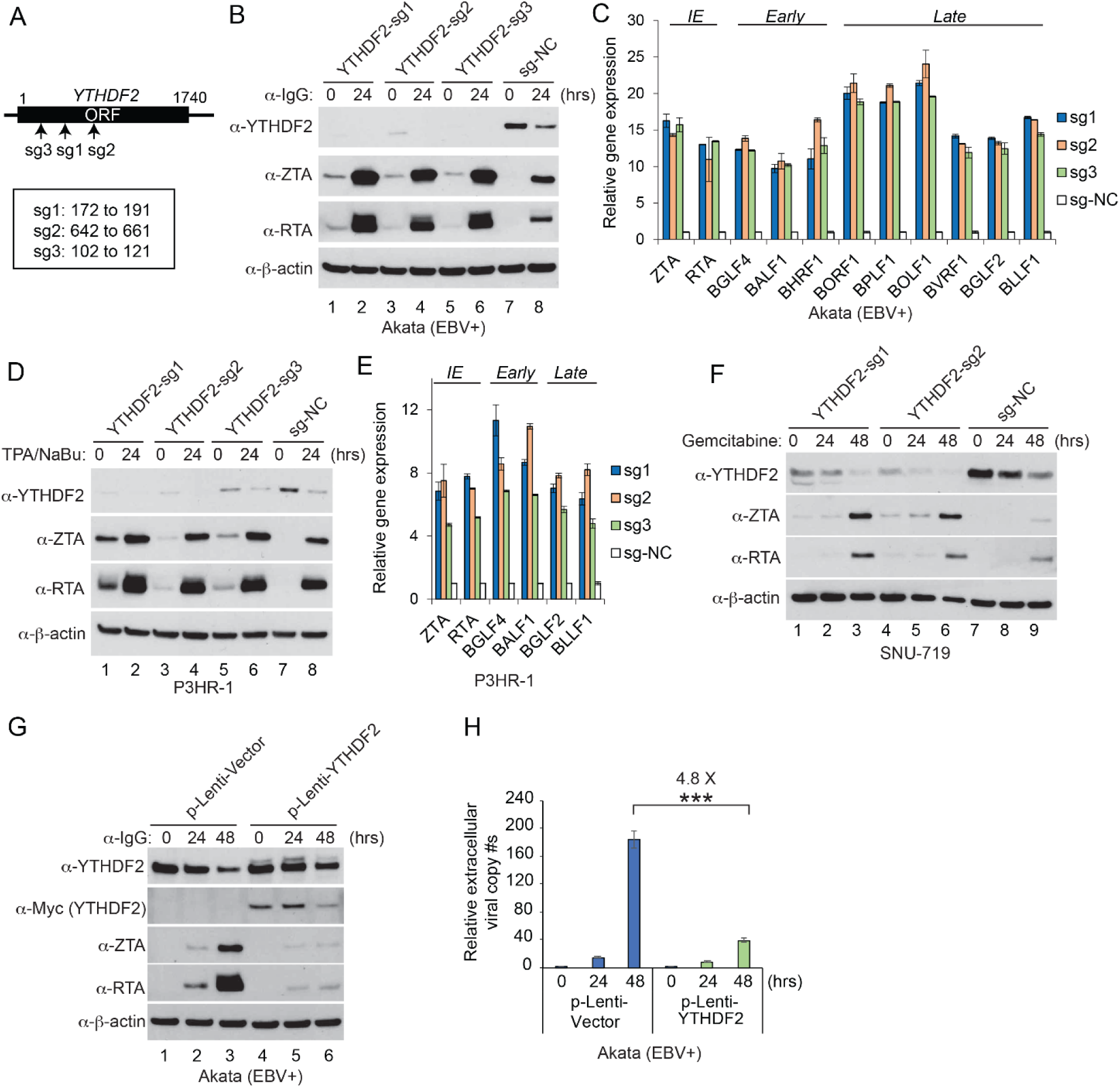
YTHDF2 restricts EBV reactivation. (A) Schematic representation showing the relative positions of Cas9 target sites for small guide RNAs sg-1 to sg-3. (B) Akata (EBV+) cells were used to establish stable cell lines using 3 different sgRNA constructs and a non-targeting control (sg-NC). The cells were untreated or lytically induced with anti-IgG-mediated cross-linking of BCR. YTHDF2 and viral protein (ZTA and RTA) expression levels were monitored by Western Blot using antibodies as indicated. (C) RNAs from YTHDF2-depleted and control Akata cells were extracted and analyzed by RT-qPCR. The values of control were set as 1. Error bars indicate ±SD. IE, immediate early gene; Early, early gene; Late, late gene. (D) P3HR-1 cells were used to establish stable cell lines as indicated. The cells were either untreated or treated with TPA and sodium butyrate (NaBu) to induce lytic reactivation. YTHDF2 and viral protein expression levels were monitored by Western Blot using antibodies as indicated. (E) RNAs from YTHDF2-depleted and control P3HR-1 cells were extracted and analyzed by RT-qPCR. The values of control were set as 1. Error bars indicate ±SD. IE, immediate early gene; Early, early gene; Late, late gene. (F) SUN-719 cells were used to establish stable cell lines as indicated. The cells were either untreated or treated with Gemcitabine to induce lytic reactivation. YTHDF2 and viral protein expression levels were monitored by Western Blot using antibodies as indicated. (G) Akata (EBV+) cells were used to establish control and YTHDF2 overexpression cell line as indicated. The cells were untreated or lytically induced by anti-IgG treatment. The expression of YTHDF2 as monitored by anti-YTHDF2 and anti-Myc antibodies. Viral protein expression levels were monitored by Western Blot using antibodies as indicated. (H) Extracellular virion-associated DNA from cells treated in panel G was extracted and the relative EBV viral copy numbers were calculated by q-PCR analysis using primers specific to BALF5. The value of vector control at 0 hr was set as 1. Results from three biological replicates are presented. Error bars indicate ±SD. ***, p<0.001. See also Figures S1-S3.

In addition, we demonstrated that YTHDF2 depletion promoted the expression of immediate-early, early and late genes even without lytic induction (**Figure 2C**), suggesting that YTHDF2 serves as a restriction factor against EBV reactivation in normal conditions. In addition, we also monitored viral latent gene expression and found that YTHDF2 depletion also enhanced the expression of EBNA1, EBNA2, EBNA3A/3B/3C, LMP1 and LMP2 (Figure S2A). Due to the key role of m^6^A RNA modification pathway in viral life cycle, we selected the m^6^A reader YTHDF2 as a starting point to explore the regulation of this pathway by caspases in EBV replication.

### YTHDF2 universally restricts EBV replication

YTHDF2 has been shown to promote or suppress KSHV lytic replication in a cell type-dependent manner, but mainly inhibit EBV replication in Burkitt lymphoma cells (25, 28, 29) (**Figure 2B** **&** 2C). To investigate whether YTHDF2 has a unified function in EBV reactivation regardless of cell type and lytic induction methods, we knocked out YTHDF2 in another Burkitt lymphoma cell line P3HR-1 and a gastric cancer cell line SNU-719, respectively. We found that YTHDF2 depletion promoted EBV gene expression in both cell lines using two different lytic inducers (**Figure 2D-2F and** **Figure S2B**), demonstrating a universal role of YTHDF2 in suppressing EBV gene expression. To determine whether YTHDF2 plays a role in EBV life cycle, we monitored the virion released to the medium upon lytic induction. We observed that YTHDF2 depletion significantly enhanced EBV copy numbers in Akata (EBV+), P3HR-1, and SNU-719 cells upon lytic induction (**Figure S3**), suggesting YTHDF2 restricts EBV replication regardless of cell type.

To further confirm the role of YTHDF2 in viral replication, we transduced the Akata (EBV+) cell with lentiviruses carrying control or the *YTHDF2* gene. YTHDF2 overexpression strongly inhibited EBV ZTA and RTA expression upon lytic induction (**Figure 2G**), and consequently abrogated the production of EBV progeny released to the medium (**Figure 2H**).

### YTHDF2 is cleaved by caspase-3, -6 and -8 at two evolutionarily conserved sites

Having shown YTHDF2 as a caspase substrate during the course of EBV replication (**Figure 1B** **&1C**), we further observed a downregulation of YTHDF2 in Akata-4E3 (EBV-) cells upon IgG cross-linking, suggesting this physiologically relevant lytic trigger plays a major role in YTHDF2 destabilization (**Figure 3A**, YTHDF2 shorter exposure, lanes 4-6). In addition to the reduction of protein level, two putative cleaved fragments of YTHDF2 were observed, whose molecular weights (MWs) are 25 and 50 kDa, respectively (**Figure 3A**, YTHDF2 longer exposure).

**Figure 3.**
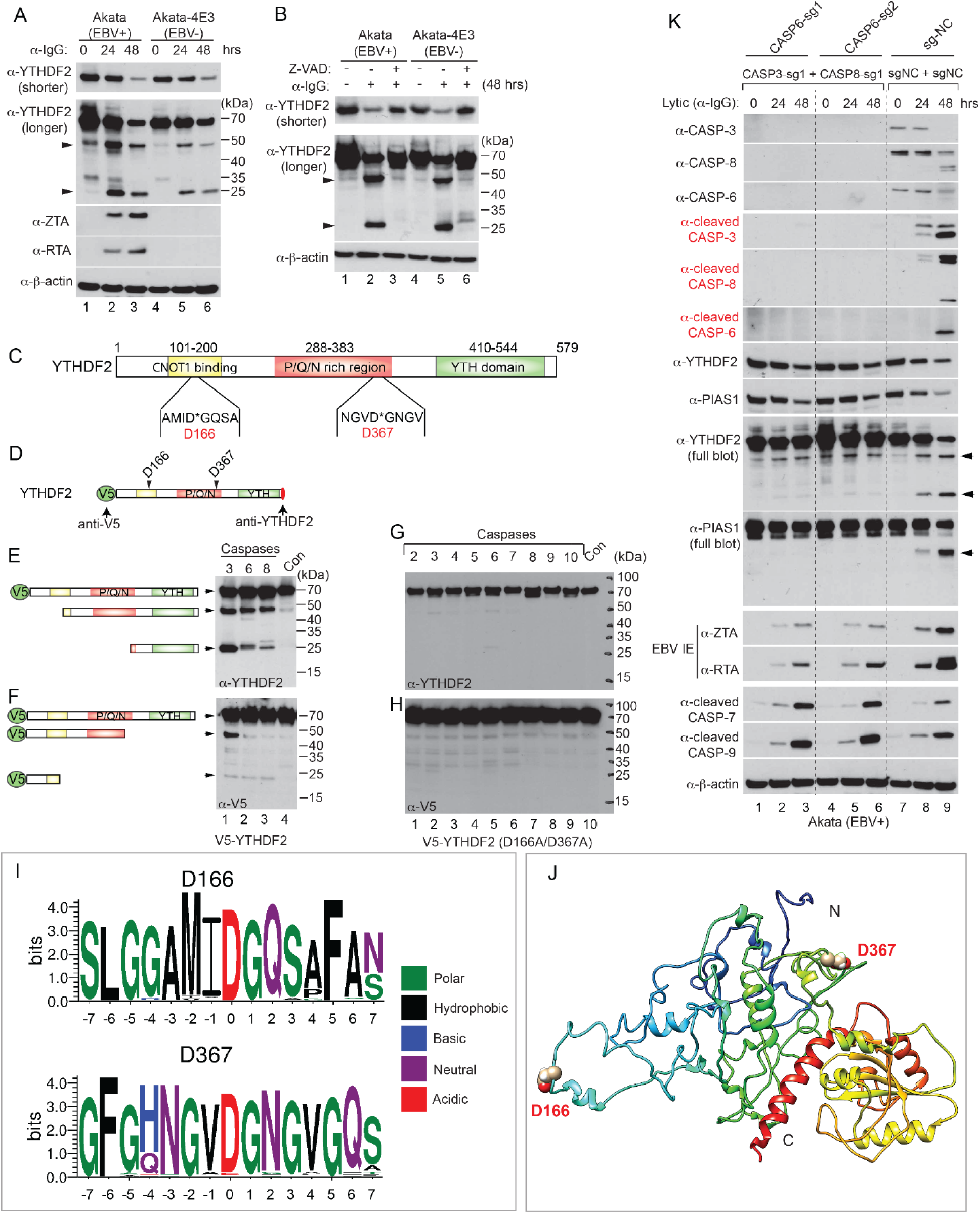
YTHDF2 is cleaved by caspases *in vivo* and *in vitro*. (A) Western Blot showing YTHDF2 downregulation by IgG cross-linking induced BCR activation. Akata (EBV+) and Akata-4E3 (EBV-) cells were treated with anti-IgG antibody as indicated. YTHDF2 and viral protein expression levels were monitored by Western Blot. Arrowheads denote cleaved YTHDF2 in the longer exposure blot. (B) Caspase inhibition blocks YTHDF2 degradation. The cells were either untreated or pretreated with a pan-caspase inhibitor (Z-VAD-FMK, 50 μM) for 1 hr, and then anti-IgG antibody was added for 48 hrs. Arrowheads denote cleaved YTHDF2. (C) Functional domains and putative cleavage sites in YTHDF2. CaspDB was used to predict the potential cleavage sites in YTHDF2. The locations of the putative cleavage sites D166 and D367 were labeled as indicated. CNOT1 binding domain: responsible for the degradation of associated RNA; P/Q/N rich region: aggregation-prone region; YTH domain: responsible for binding to m^6^A-modified RNA. (D) Schematic representation of V5-tagged YTHDF2 with two putative cleavage sites. Red oval, anti-YTHDF2 monoclonal antibody recognition site. (E-F). Wild-type V5-YTHDF2 was incubated with individual recombinant caspase for 2 hrs. Western Blot was performed using either anti-YTHDF2 (E) or anti-V5 (F) antibodies. The relative position of predicted cleavage fragments was labeled as indicated. (G-H) YTHDF2 (D166A/D367A) mutant protein was incubated with individual recombinant caspase for 2 hrs. Western Blot was performed using antibodies as indicated. (I) Motif analysis showing the conservation of the two cleavage sites and the surrounding amino acids. Amino acid sequences were extracted from 97 (D166) and 80 (D367) vertebrate species and motif logos were generated using WebLogo. (J) Structure modeling of full-length YTHDF2 by I-TASSER. The two cleavage sites D166 and D367 are labeled as indicated. N and C denote N-terminus and C-terminus, respectively. (K) Triple depletion of caspase-3, -8 and -6 reduces YTHDF2 and PIAS1 degradation and blocks viral protein accumulation. The CASP3/CASP8/CASP6-triply-depleted Akata (EBV+) cells were lytically induced by anti-IgG treatment. The expression of caspases, cleaved caspases, YTHDF2, PIAS1 and viral proteins (ZTA and RTA) was monitored by Western Blot using antibodies as indicated. Arrowheads denote cleaved fragments. See also Figures S4-S8 and Table S2.

As expected, caspase inhibition also blocked the generation of two fragments in both Akata (EBV+) and Akata-4E3 (EBV-) cells (**Figure 3B**, YTHDF2 longer exposure). We predicted that the 25 kDa fragment likely represents a C-terminal cleaved YTHDF2 (aa 368-579, predicted MW=24 kDa). Similarly, YTHDF2 was also downregulated in lytically induced P3HR-1 (**Figure S4A, lanes 1-2**) and SNU-719 cells (**Figure S4B**, lanes 1-2). Caspase inhibition blocked YTHDF2 degradation and inhibited viral replication (**Figure S4A-S4B**, lanes 3-4) (7).

To determine how caspase activation and YTHDF2 degradation are associated with EBV reactivation, we monitored the activation of caspase, YTHDF2 and EBV ZTA/RTA protein levels at various time points after lytic induction. We found that the activation of caspase correlated well with YTHDF2 degradation and the expression of EBV ZTA and RTA in Akata (EBV+), P3HR-1, and SNU-719 cells (**Figure S5A-S5C**). To determine whether other apoptotic inducer could induce EBV reactivation, we used Taxol to induced intrinsic apoptotic pathway and found that this treatment promoted caspase activation, YTHDF2 degradation and EBV ZTA and RTA expression in all three cell lines tested (**Figure S5D-S5F**).

To further demonstrated the role of caspase activation in EBV replication, we overexpressed caspase-8 in Akata (EBV+) cells using lentiviral transduction. We found that caspase-8 overexpression promoted EBV ZTA and RTA protein accumulation upon lytic induction (**Figure S6A**), and consequently fostered the production of EBV progeny released to the medium (**Figure S6B**). Caspase-8 overexpression not only promoted lytic gene expression but also facilitated latent gene expression upon lytic induction (**Figure S6C**).

To determine the major caspases responsible for YTHDF2 cleavage, we performed an in vitro cleavage assay using individual recombinant caspases and YTHDF2. We found that caspase-3/-6/-8, and, to a lesser extent, caspase-7 all cleaved YTHDF2 (**Figure S4C**). The sizes of cleaved fragments *in vitro* were similar to those observed in cells, suggesting that YTHDF2 is a *bona fide* caspase substrate *in vitro* and *in vivo*.

In addition to the predicted cleavage sites D367 (**Figure 1A**), we used an *in silico* prediction algorithm (CaspDB) to predict additional cleavage sites (39). A second highest scoring cleavage site (D166) was identified near the N-terminal of YTHDF2 (**Figure 3C**).

To explore whether there are two cleavage sites in YTHDF2, we first generated an N-terminal V5-tagged YTHDF2 by a method we used previously for PIAS1 (7), which allows the determination of both N- and C-terminal cleavage fragments by anti-V5 and anti-YTHDF2 (C-terminal) monoclonal antibodies, respectively (**Figure 3D**). Given the sizes of these cleaved bands recognized by the C-terminal YTHDF2 antibody, we noticed that caspase-3, -6 and -8 mediated cleavage led to one ∼50 kDa C-terminal fragment, and caspase-3, and to a lesser extend caspase-6 and -8, generated one 25 kDa C-terminal fragment (**Figure 3E**). The anti-V5 antibody revealed one ∼50 kDa N-terminal cleavage fragment generated by caspase-3 and one smaller N-terminal (slightly less than 25 kDa) fragment generated by all 3 caspases (**Figure 3F**). Together, these results demonstrate that YTHDF2 is indeed cleaved at two sites by caspase-3, -6 and -8.

To further confirm whether D166 and D367 are the major cleavage sites, we mutated these two residues to alanines (D166A/D367A) and then examined the cleavage profile using in vitro cleavage assays (**Figure 3G** **& 3H**). Consistent with our prediction, mutations on those two sites prevented YTHDF2 cleavage by caspases, indicating that D166 and D367 are the major cleavage sites. To examine the conservation across different species of these two cleavage motifs (AMID*G and NGVD*G) within YTHDF2, we extracted 15 amino acids surrounding the cleavage sites from 80-97 vertebrate species. Interestingly, we found that not only the cleavage motifs but also the surrounding amino acids are highly conserved (**Figure 3I** **and** **Figure S4D**), suggesting a key regulatory function for YTHDF2 cleavage during evolution.

Considering cleavage sites are normally exposed to the surface of the protein, we used an I-TASSER (Iterative Threading ASSEmbly Refinement) algorithm (40, 41) to generate a 3-dimensional (3D) structure for full-length YTHDF2 by integrating the crystal structure of the YTH domain (42, 43). One structure with two cleavage sites located on the protein surface was visualized with Chimera (44) and both sites fall within flexible regions favoring cleavage by caspases (**Figure 3J**).

Because caspase-3, -6 and -8 are the major caspases that can redundantly cleave YTHDF2 (**Figure S4C**) and another restriction factor PIAS1 (7), we created two *CASP3/CASP8/CASP6* (genes encoding caspase-3, -8 and -6) triply depleted Akata (EBV+) cell lines by CRISPR/Cas9 approaches (**Figure 3K**; total and cleaved caspase-3, -8 and -6 blots). The depletion of these three caspases alleviated the degradation of YTHDF2 and PIAS1 upon lytic induction (**Figure 3K**; YTHDF2 and PIAS1 blots, lanes 2-3, 5-6 vs 8-9). Consequently, the gene expression and protein accumulation of EBV ZTA and RTA was also significantly reduced in *CASP3/CASP8/CASP6-*depleted cells (**Figure S7A-S7B** and **Figure 3K**; ZTA and RTA blots, lanes 2-3, 5-6 vs 8-9). As expected, caspases depletion also suppressed the production EBV progeny released from the cells (**Figure S7C**).

Interestingly, the depletion of caspase-3, -6 and -8 led to a slight increase of caspase-7 and -9 activation, which may contribute to the partial destabilization of YTHDF2 and PIAS1 upon lytic induction (**Figure 3K**; YTHDF2 and PIAS1 blots, lanes 1 vs 3 and 4 vs 6).

To further demonstrate whether apoptotic induction is associated with YTHDF2 degradation and EBV lytic protein expression in single cell level, we performed immunofluorescence assays. We found that lytic induction gradually led to enhanced uptake of propidium iodide (PI), a marker of late apoptosis, and reduced YTHDF2 signals in Akata (EBV+) cells (**Figure 4A**). Consistently, we found that reduced YTHDF2 level correlated well with enhanced EBV EAD expression in single cell level (**Figure 4B**). Similarly, lytic induction also triggered the uptake of PI, reduction of YTHDF2, and enhancement of EBV EAD expression in SNU-719 cells (**Figure 4C and 4D**).

**Figure 4.**
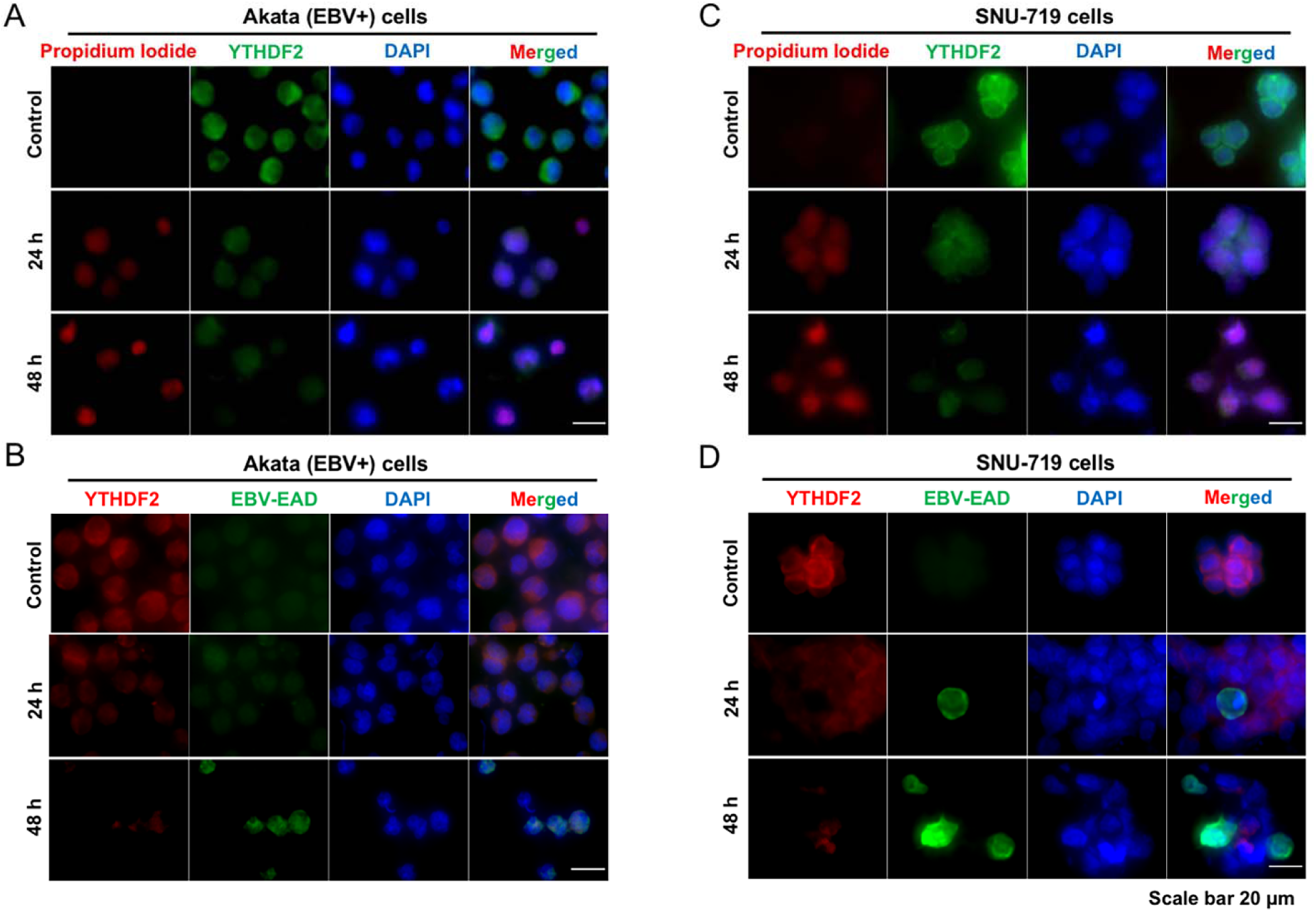
Apoptotic induction triggers YTHDF2 degradation and EBV lytic protein expression. (A-B) Immunofluorescence assay showing YTHDF2 downregulation and EBV EAD upregulation in apoptotic Akata (EBV+) cells upon lytic induction. Akata (EBV+) cells were either untreated (control) or treated with anti-IgG antibody for 24 and 48 hrs as indicated. (A) The cells were stained with Propidium Iodide (PI) and then permeabilized for immnunostaining with anti-YTHDF2 antibody. (B) Cells were permeabilized and co-immunostained with anti-YTHDF2 and anti-EBV EAD antibodies as indicated. (C-D) Immunofluorescence assay showing YTHDF2 downregulation and EBV EAD upregulation in apoptotic SNU-719 cells upon lytic induction. SNU-719 cells were either untreated (control) or treated with TPA and NaBu for 24 and 48 hrs as indicated. (C) Cells were stained with Propidium Iodide (PI) and then permeabilized for immunostaining with anti-YTHDF2 antibody. (D) Cells were permeabilized and co-immunostained with anti-YTHDF2 and anti-EBV EAD antibodies as indicated. Scale bar, 20 μm

Because caspases, especially caspase-8, have been shown to promote apoptosis but suppress necroptosis (45, 46), we also tested whether caspase inhibition could enhance the necroptotic pathway upon lytic induction. We monitored the level of necroptotic markers (phospho-RIP and phospho-RIP3) following lytic induction or the combination of caspase inhibition and lytic induction. We found that lytic induction led to reduced levels of phospho-RIP and phospho-RIP3 (**Figure S8A and S8B,** lane 1 vs 2) while the combination of caspase inhibition and lytic induction did not enhance the levels of phospho-RIP and phospho-RIP3 in both Akata (EBV+) and P3HR-1 cells (**Figure S8A** **and** **S8B**, lane 2 vs 3 and 4). In SNU-719 cells, both lytic induction and caspase inhibition plus lytic induction did not affect the levels of phospho-RIP and phospho-RIP3 (**Figure S8C**). These results suggested that caspase inhibition does not further affect the necroptotic pathway when cells were treated with EBV lytic-inducing agents.

These results suggested that lytic induction induced caspase activation and the subsequent cleavage of host restriction factors, YTHDF2 and PIAS1, promote EBV lytic replication (7).

### Caspase-mediated YTHDF2 cleavage promotes EBV replication

To further investigate whether caspase-mediated degradation of full length YTHDF2 abrogates its restriction toward EBV lytic replication, we first generated a CRISPR-resistant YTHDF2 variant in a lentiviral vector. This was achieved by introducing a silent mutation on the YTHDF2-sg2 PAM sequence (**Figure 5A**) (7, 38). We first transduced the Akata (EBV+) cells with lentiviruses carrying vector control, WT and mutant (D166A/D367A) YTHDF2 to establish cell lines. We then depleted endogenous YTHDF2 by lentiviral transduction of YTHDF2-sg2 into these three cell lines. We then measured YTHDF2 protein degradation, viral protein accumulation and viral DNA replication after lytic induction. YTHDF2 (D166A/D367A) mutant was resistant to caspase-mediated cleavage and therefore became more stable than the WT counterpart upon lytic induction (**Figure 5B**, lane 6 vs 9). Compared to WT YTHDF2, the D166A/D367A mutant strongly suppressed EBV protein accumulation (**Figure 5B**), and consequently reduced the intracellular and extracellular viral copy numbers upon lytic induction (**Figure 5C**), further consolidating caspase-mediated cleavage in antagonizing the anti-viral function of YTHDF2.

Caspase-mediated cleavage led to not only a decrease of total YTHDF2 protein level but also an increase of cleaved fragments (**Figure 3A** **& 3B**). We hypothesized that these cleavage fragments compete with WT YTHDF2 to promote viral replication. To test this hypothesis, we created a series of cleavage-mimicking fragments using a lentiviral vector (**Figure 5D**). We then transduced SNU-719 cells to establish individual cell lines. All these fragments promoted the protein and mRNA expression of EBV ZTA and RTA upon lytic induction in these gastric cancer cell lines (**Figure 5E**, lane 1 vs 2-6; **Figure S9A**). To further confirm these results, we established fragment-expressing cell lines using Akata (EBV+) cells. F2 (aa 1-367)-expressing cell line failed being established after multiple attempts. However, in other four cell lines established, these cleavage-mimicking fragments strongly promoted EBV ZTA/RTA protein and *ZTA/RTA* mRNA expression (**Figure 5F**, lane 1 vs 2-5 and lane 6 vs 7-10; **Figure S9B**). To further determine whether these results are due to protein overexpression or not, we created a pLenti-Halo control and transduced Akata (EBV+) cells to establish a Halo-expressing cell line. We also included vector control, WT YTHDF2, and F1 (aa 1-166)/F4 (aa 167-579)-expressing cell lines for comparison. We found that Halo expression had similar effects as vector control in EBV ZTA and RTA protein expression (**Figure S9C**, lane 1 vs 2 and lane 6 vs 7), and that WT YTHDF2 suppressed but F1 (aa 1-166) and F4 (aa 167-579) promoted the expression of EBV ZTA and RTA upon lytic induction (**Figure S9C**, lanes 1-2 vs 3-5 and lanes 6-7 vs 8-10). As expected, we observed that viral RNA level correlated well with viral protein expression in these cell lines (**Figure S9D**).

**Figure 5.**
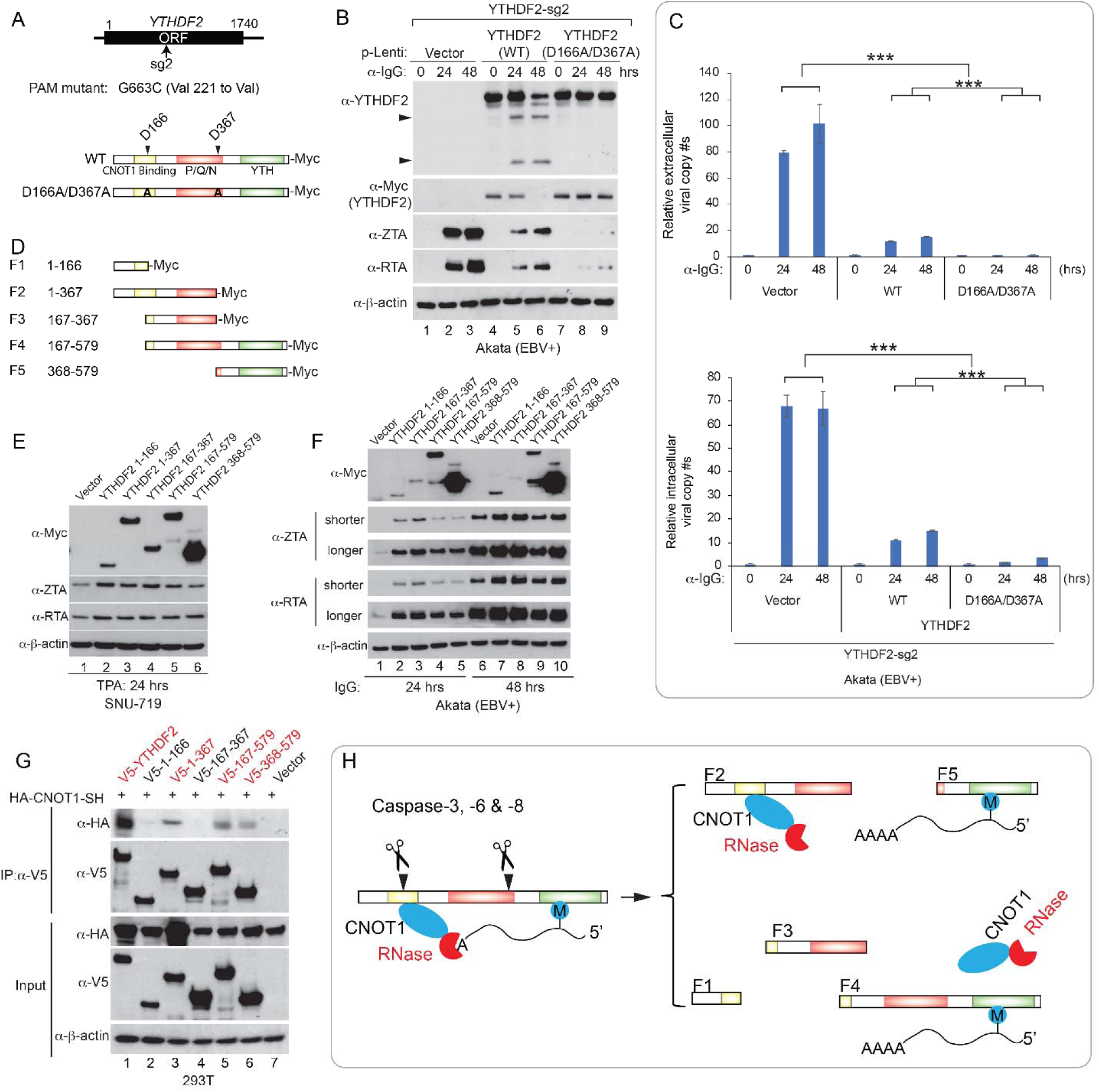
YTHDF2 cleavage promotes EBV replication. (A) The design of CRISPR/Cas9-resistant YTHDF2 variant was based on the sg-2 protospacer adjacent motif (PAM). D166A/D367A mutations were introduced into the PAM-mutated YTHDF2. Both constructs were cloned into a lentiviral vector with a C-terminal Myc-tag. (B-C) WT and cleavage-resistant YTHDF2 suppresses EBV replication. Akata (EBV+) YTHDF2-sg2 cells were reconstituted with WT or cleavage-resistant YTHDF2 (D166A/D367A) using lentiviral constructs. Western Blot analysis showing YTHDF2 and EBV protein expression levels in these cell lines upon IgG cross-linking as indicated (B). Arrowheads denote cleaved fragments. Extracellular and intracellular viral DNA was measured by qPCR using primers specific to BALF5 (C). The value of vector control at 0 hr was set as 1. Results from three biological replicates are presented. Error bars indicate ±SD. **, p<0.01; ***, p<0.001. (D) Schematic representation of 5 YTHDF2 cleavage-mimicking fragments. These fragments were cloned into a lentiviral vector with a C-terminal Myc-tag. (E) SNU-719 cells were transduced with lentiviruses carrying vector control or individual fragment to establish stable cell lines. Western Blot analysis showing YTHDF2 fragments and EBV protein expression levels in these cell lines upon lytic induction by adding TPA (20 ng/ml) for 24 hrs. (F) Akata (EBV+) cells were transduced with lentiviruses carrying vector control or individual fragment to establish stable cell lines. Western Blot analysis showing YTHDF2 fragments and EBV protein expression levels in these cell lines upon lytic induction by anti-IgG treatment for 24 and 48 hrs. Shorter and longer exposures were included to show the differences in protein levels. (G) Caspase-mediated cleavage impairs YTHDF2 binding to CNOT1. Halo-V5-tagged WT YTHDF2 and the individual fragments were co-transfected with HA-tagged CNOT1 SH domain into 293T cells as indicated. Co-immunoprecipitation (Co-IP) experiments were performed using anti-V5 antibody-conjugated magnetic beads. The immunoprecipitated samples and total cell lysates (Input) were analyzed by Western Blot with antibodies as indicated. (H) Model showing the functional consequences of YTHDF2 cleavage in CNOT1 binding and the targeting of m^6^A-modified RNA. See also Figures S9 and S11.

YTHDF2 has been shown to bind CNOT1 to recruit CCR4-NOT deadenylase complex for RNA decay (18). To determine whether YTHDF2 cleavage impairs its binding to CNOT1, we co-transfected a CNOT1-SH domain-expressing construct with individual YTHDF2 fragment-expressing vector into 293T cells and then performed a co-immunoprecipitation (co-IP) assay. As expected, we found that CNOT1 is co-IPed with WT YTHDF2 (**Figure 5G**, lane 1). We also observed weaker binding signals of CNOT1 with F2 (aa 1-367), F4 (aa 167-579) and F5 (aa 368-579) fragments, compared to full-length YTHDF2 (**Figure 5G**, lane 1 vs 3, 5 & 6). This is in part consistent with a previous study showing that YTHDF2 (aa 100-200) binds to CNOT1 (18). The binding between CNOT1 and F4 (aa 167-579)/F5 (aa 368-579) fragments could be mediated by a bridge protein or RNA. However, our study indicated that the YTH domain coordinates with the N-terminal region for enhanced YTHDF2 binding to CNOT1. Interestingly, two fragments, F1 (aa 1-166) and F3 (aa 167-367), lost the interaction with CNOT1, suggesting an essential N-terminal binding site for CNOT1 locating near the caspase cleavage site D166 of YTHDF2 (**Figure 5G**, lane 3 vs 2 & 4). Together, these results suggested that caspase mediated cleavage could convert YTHDF2 from an anti-viral restriction factor to several pro-viral fragments (**Figure 5H**).

### YTHDF2 regulates viral and cellular gene stability to promote viral replication

It is known that YTHDF2 binds to m^6^A-modified viral and cellular RNAs to control their stability (11, 16, 29, 47). By m^6^A RNA immunoprecipitation (RIP) followed by reverse transcription quantitative real-time PCR (RT-qPCR) analysis, we found that EBV immediate early (*ZTA/BZLF1* and *RTA/BRLF1*) and early (*BGLF4*) transcripts are modified by m^6^A, not only revealing a new m^6^A target *BGLF4* but also confirming the results for *ZTA and RTA* captured by m^6^A-sequencing (m^6^A-seq) analyses (25) (**Figure S10A**). We further demonstrated that YTHDF2 strongly binds to *ZTA, RTA* and *BGLF4* transcripts given the results from YTHDF2 RIP coupled with RT-qPCR analysis (**Figure S10B**). Together, all these results suggested that YTHDF2 binds to EBV lytic transcripts through m^6^A modifications (**Figure S10**) and then promotes their decay to restrict EBV reactivation (**Figure 2C** **& 2E**).

In addition to regulating viral mRNA stability, YTHDF2 was reported to modulate m^6^A-modified cellular transcripts to affect a variety of cellular processes. YTHDF2 targets were identified by YTHDF2 RIP and PAR-CLIP (photoactivatable ribounucleoside crosslinking and immunoprecipitation) datasets from previous publications (16, 48). We found a group of YTHDF2 target genes involved in the biological process of “activation of cysteine-type endopeptidase activity involved in apoptotic process”, also known as “caspase activation” (**Figure S11A**). Majority of these proteins have protein-protein interactions with caspase-8 (CASP8). As caspase activation plays a critical role in promoting EBV reactivation (**Figure 3K**) (7), we reasoned that YTHDF2 may regulate these genes to foster viral replication. The RT-qPCR results revealed that the mRNA levels of many potential YTHDF2 targets were elevated after YTHDF2 was knocked out by CRISPR/Cas9. Particularly, *CASP8* (encoding caspase-8) had more than 2-fold significant increase in both Akata (EBV+) and P3HR-1 cells (**Figure 6A** **& 6B**).

**Figure 6.**
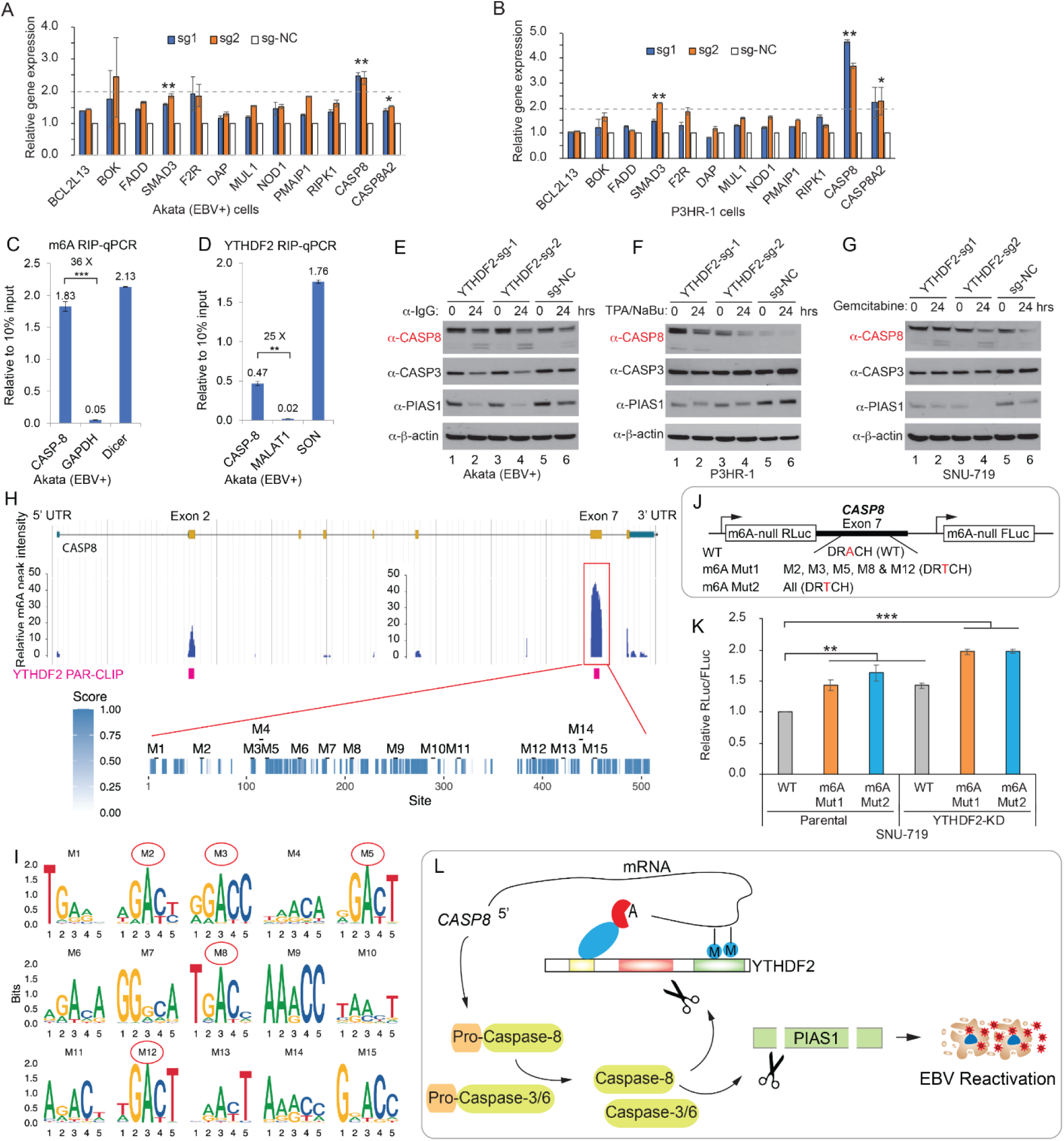
YTHDF2 regulates *CASP8* mRNA stability through m^6^A modifications. (A-B) YTHDF2 depletion promotes *CASP8* mRNA expression. Akata (EBV+) cells and P3HR-1 cells carrying different sgRNA targeting YTHDF2 or control (sg-NC) were used to extract total RNA and qPCR analyses were performed a group of YTHDF2-targeted cellular genes involved in caspase activation. The values were normalized with a non YTHDF2 target *HPRT1*. The values of sg-NC were set as 1. (C-D) *CASP8* is modified by m^6^A and YTHDF2 binding to *CASP8*. Akata (EBV+) cells were used to perform m6A RIP-qPCR (C) and YTHDF2 RIP-qPCR (D), respectively. Values are displayed as fold change over 10% input. (E-G) YTHDF2 depletion promotes caspase-8 protein expression and PIAS1 cleavage upon lytic induction. Akata (EBV+) cells (E), P3HR-1 cells (F) and SNU-719 cells (G) carrying different sgRNA targeting YTHDF2 or control (sg-NC) were lytically induced by anti-IgG, TPA and sodium butyrate (NaBu) and gemcitabine treatment for 24 hrs. Protein expression was monitored by Western Blot using antibodies as indicated. (H) *CASP8* m^6^A peaks were extracted from MeT-DB V2.0 database. YTHDF2-PAR-CLIP data were retrieved from Wang et al.(16). The Exon-7 of *CASP8* with highest m^6^A peaks were analyzed for conservation among sequences derived from 100 vertebrate species. 15 potential m^6^A motifs (M1-M15) were extracted based on m^6^A motif DRACH. (I) Motif logos were generated for 15 individual sites. Red cycles denote highly conserved motifs (M2, M3, M5, M8 and M12) across 100 vertebrate species. (J-K) WT and mutant *CASP8*-Exon-7 were cloned into the m^6^A-null Renilla luciferase (RLuc) reporter (3’UTR region) that also express Firefly luciferase (FLuc) from a separate promoter (J). These three reporter plasmids were transfected into parental or YTHDF2-depleted (YTHDF2 KD) SNU719 cells. Relative Renilla to Filefly luciferase activity (RLuc/FLuc) was calculated (K). The value of WT in parental cells was set as 1. (L) Model illustrating YTHDF2 regulation of *CASP8* mRNA and caspase-8 regulation of YTHDF2 and PIAS1 in EBV reactivation. Results from three biological replicates are presented. Error bars indicate ±SD. *, p< 0.05; **, p< 0.01; ***, p< 0.001. See also Figures S10, S11 and Table S3.

To test whether *CASP8* is modified by m^6^A in EBV-positive cells, we performed m^6^A RIP followed by RT-qPCR using Akata (EBV+) cells. The results demonstrated that *CASP8* mRNA is strongly modified by m^6^A, compared to the controls (**Figure 6C**). In addition, YTHDF2 indeed bound to *CASP8* mRNA as revealed YTHDF2 RIP-qPCR analysis (**Figure 6D**). Consistent with mRNA elevation, caspase-8 protein level was also increased upon YTHDF2 depletion (**Figure 6E-6G** lanes 1, 3 vs 5). Upon lytic induction, we observed enhanced caspase-8 activation and consequently PIAS1 degradation (**Figure 6E-6G**, lanes 2, 4 vs 6). Conversely, WT and D166A/D367A mutant YTHDF2 reconstitutions led to reduced caspase-8 protein level and caspase activation upon lytic induction, with D166A/D367A mutant having the strongest effects (**Figure S11B**, lanes 1-3 vs 4-6 & 7-9). By RT-qPCR analysis, we further found that *CASP8* mRNA level was increased in cells expression WT YTHDF2 but not the D166A/D367A mutant following lytic induction (**Figure S11C**). To understand the relationship between YTHDF2 regulation of caspase-8 and EBV replication, we pre-treated the YTHDF2-depleted Akata (EBV+) cells with or without a caspase-8 inhibitor and then induced the lytic cycle by anti-IgG treatment. We found that caspase-8 inhibition reduced EBV ZTA and RTA expression and consequently inhibited EBV virion production even when YTHDF2 is depleted (**Figure S11D** **& S11E**, lane 7 vs 9, lane 8 vs 10, lane 12 vs 14 & lane 13 vs 15). To further confirm our results, we created a *CASP8/YTHDF2*-double knockout cell line using Akata (EBV+) cells. We found that, compared with *YTHDF2*-knockout cells, both EBV lytic protein expression and viral copy numbers were significantly reduced in *CASP8/YTHDF2*-double knockout cells (**Figure S11F**, lanes 2-3 vs 5-6 and **Figure S11G**).

Together, our results suggested that the depletion of YTHDF2 can promote EBV reactivation partially through enhanced capase-8 activation.

M^6^A modifications are mainly located near the stop codon region to regulate mRNA decay (16, 49, 50). By mining the m^6^A modification database (51, 52) and YTHDF2-PAR-CLIP datasets (16), we found that *CASP8-*Exon-7 has the highest m^6^A peaks that are closed to the stop codon (**Figure 6H**). Using conserved m^6^A motif DRACH (D=G/A/U, R=G/A, H=A/U/C) (53), we identified 15 potential m^6^A sites that are evenly distributed across the entire Exon-7 (**Figure 6H**). We found that Motif-2 (M2), M3, M5, M8 and M12 are highly conserved across more than 90 vertebrates ranging from human, mouse, bat to zebrafish, while the others likely evolve gradually during evolution (**Figure 6I** **and Table S3**).

To determine whether m^6^A modifications contribute to *CASP8* mRNA stability, we synthesized the Exon-7 DNA of WT *CASP8* [without the first 20 base bases (bps)], and its mutant counterparts with conserved motifs (M2, M3, M5, M8 and M12) or all putative motifs mutated (**Figure 6J**). Luciferase reporters were created with WT or mutant DNA inserted into the 3’-untranslated region (3’-UTR) of an m^6^A-null vector (**Figure 6J**). These reporters were modified to silently mutate all putative m^6^A sites in the *Renillia* and *Firefly luciferase* genes to specifically test the function of m^6^A modifications on *CASP8-*Exon-7 (54). We then transfected these plasmids individually into SNU-719 cells carrying YTHDF2 or with YTHDF2 depleted by CRISPR/Cas9. Disruption of m^6^A modification by mutations enhanced the relative luciferase activity compared to the WT reporter (**Figure 6K**). In addition, depletion of YTHDF2 also enhanced the relative luciferase activity (**Figure 6K**), suggesting that YTHDF2 directly regulates *CASP8* stability. We also noticed a further enhancement of relative luciferase activity for mutant reporters in YTHDF2 depleted cells (**Figure 6K**), suggesting a regulation of *CASP8* stability by additional m^6^A readers, e.g. YTHDF1 or YTHDF3 (20).

Together, these results suggested that YTHDF2 depletion by CRISPR/Cas9 or its cleavage by caspases could promote viral reactivation through up-regulating caspase-8 which further enhances the cleavage of antiviral restriction factors, including PIAS1 and YTHDF2 (**Figure 6L** **and Figure S11B & S11C**) (7).

### M^6^A pathway proteins are regulated by caspase-mediated cleavage

The m^6^A RNA modification machinery contains writers, readers and erasers that mediate methylation, RNA binding and demethylation steps, respectively (**Figure 7A**). Prompted by our YTHDF2 results, we predicted that other members involved in the m^6^A pathway may be downregulated by caspase-mediated cleavage during viral replication. To evaluate this possibility, we first monitored the protein levels of additional m^6^A readers (YTHDF1, YTHDF3, YTHDC1 and YTHDC2), m^6^A writers (METTL3, METTL14, VIRMA and WTAP), and m^6^A erasers (ALKBH5 and FTO). Interestingly, except ALKBH5, all other proteins were significantly down-regulated upon lytic induction (**Figures 7B** **&** **S12A-S12K**). As a control, the protein level of a putative DNA N6-methyladenine (6mA) writer N6AMT1 did not change upon lytic induction (**Figures 7B** **&** **S12L**), suggesting a specific regulation of the m^6^A RNA modification pathway by cellular caspases.

**Figure 7.**
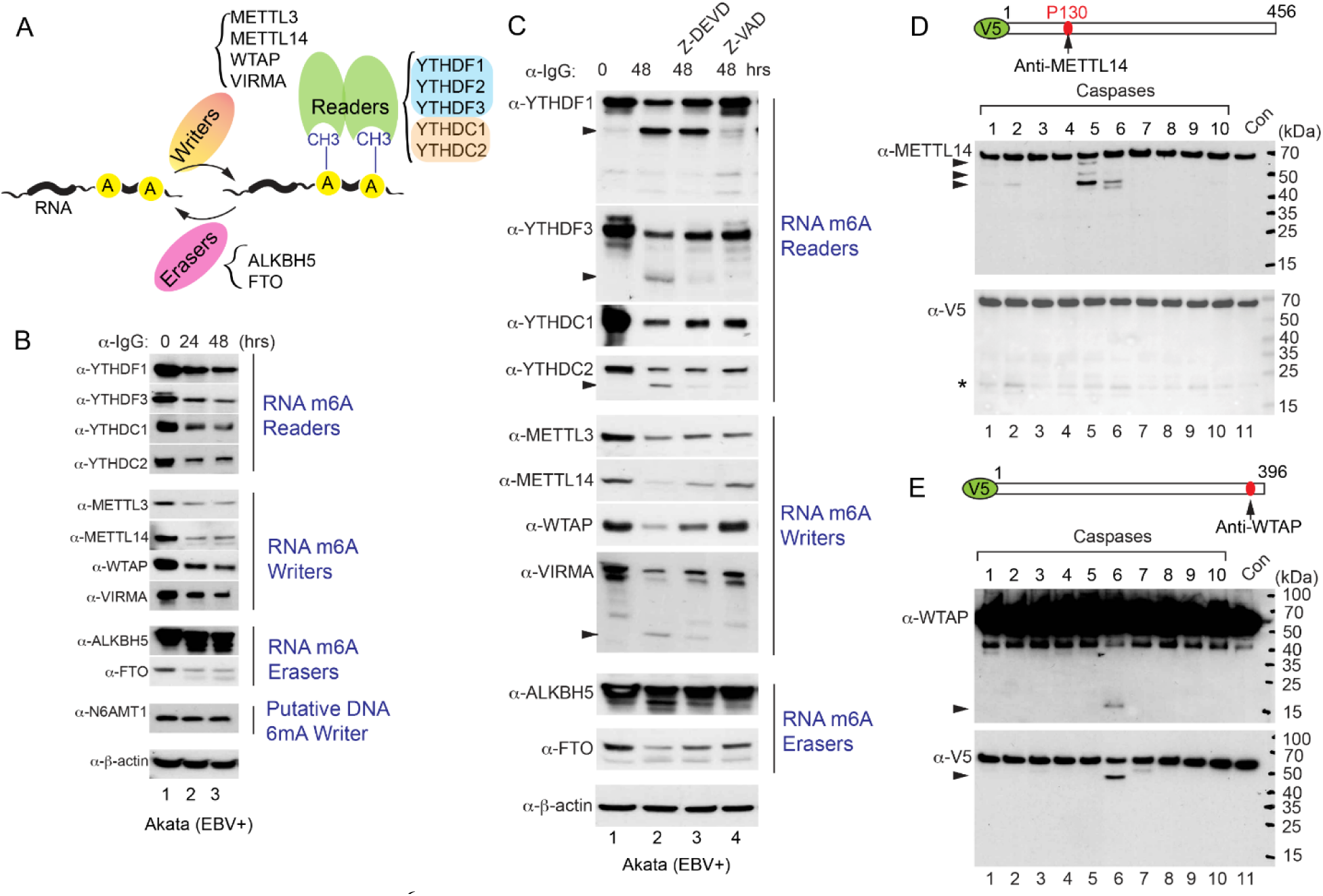
Caspases cleave m^6^A RNA modification pathway proteins. (A) Diagram summarizing the major writers, readers and erasers involved in the m^6^A RNA modification pathway. (B) The downregulation of m^6^A RNA modification pathway proteins during EBV reactivation. Akata (EBV+) cells was treated with anti-IgG antibody to induce EBV reactivation for 0, 24 and 48 hrs. Western Blot was performed using antibodies as indicated. N6AMT1 and β-actin blots were included as controls. (C) Caspase inhibition blocks the degradation of m^6^A RNA modification pathway proteins. The Akata (EBV+) cells were either untreated or pretreated with a caspase-3/-7 inhibitor (Z-DEVD-FMK, 50 μM) or pan-caspase inhibitor (Z-VAD-FMK, 50 μM) for 1 hr, and then anti-IgG antibody was added for 48 hrs. Western Blot was performed using antibodies as indicated. (D and E) V5-METTL14 (D) and V5-WTAP (E) were incubated with individual caspase for 2 hrs at 37°C. Western Blot was performed using anti-METTL14, anti-V5 and anti-WTAP antibodies as indicated. The locations of antibody recognition epitopes were labelled as indicated. Arrowheads denote cleaved fragments. Star denotes non-specific bands. See also Figures S12-S14.

To further demonstrate the role of caspases in the downregulation of m^6^A pathway proteins, we examined their protein levels in the presence of caspase-3/7 or pan-caspase inhibitors. Indeed, we found that caspase inhibition could restore the protein levels of YTHDF1, YTHDF3, METTL14, WTAP, VIRMA, and FTO (**Figure 7C**). The partial restoration of YTHDC1, YTHDC2, and METTL3 suggested that these proteins are controlled partially by caspase cleavage and partially by other protein degradation mechanisms (**Figure 7C**).

YTHDF2 shares high sequence homology with YTHDF1 and YTHDF3. Sequence alignment showed that the cleavage motif AMID*G, but not NGVD*G, is partially conserved among YTHDF family proteins (**Figure S13A**). Because TVVD*G in YTHDF1 and AITD*G in YTHDF3 are also evolutionary conserved (**Figure S13B & S13C**), we reasoned that YTHDF1 and YTHDF3 are subjected to caspase mediated cleavage on the N-terminal sites. Consistent with our prediction, not only did YTHDF1 and YTHDF3 protein levels decrease, a cleaved band near 50 kDa was generated upon lytic induction by IgG-crosslinking, matching the calculated molecular weights of C-terminal fragments generated by cleavage on the N-terminal sites (**Figure S12B & S12C**). In addition, cleaved fragments were detected for YTHDC1, YTHDC2, METTL14, VIRMA and FTO (**Figure S12D-S12K**).

To further demonstrate that the m^6^A writers are cleaved by caspase, we performed in vitro cleavage assay using purified proteins. Interestingly, we found that METTL14 is mainly cleaved by caspase-2, -5, and -6 revealed by anti-METTL14 antibody (**Figure 7D**), whereas WTAP is mainly cleaved by caspase-6 (**Figure 7E**). However, we only observed trace amount of METTL3 cleavage by caspase-4, -5, -6 and -7 revealed by anti-METTL3 antibody (**Figure S14A**).

The cleavage patterns suggested that METTL14 is cleaved on multiple sites while WTAP is possibly cleaved on one major site near the C-terminus (**Figure 7E**). Based on the C-terminal fragment molecular weight (15 kDa), we reasoned that D301 or D302 is cleaved. To examine this, we generated WTAP mutants (D301A and D301A/D302A) and performed in vitro cleavage experiments. Interestingly, only the D301A/D302A mutant could block WTAP cleavage (**Figure S14B**), indicating D302 as the major cleavage site. Sequence analysis revealed that the core cleavage motif TEDD*F is evolutionarily conserved (**Figure S14C**). Compared to cleavage sites normally followed by glycine, serine or alanine (36), the discovery of a conserved phenylalanine after the cleavage site extends our knowledge on the substrates recognition by caspase (**Figure S14D**).

### Depletion of m^6^A writers (METTL3, METTL14, WTAP or VIRMA) and reader YTHDF3 promotes EBV replication

The aforementioned studies suggest that YTHDF2 restricts viral replication through m^6^A modifications. We reasoned that the disruption of the m^6^A writer complex and other readers might also promote viral replication. To test our hypothesis, we used a CRISPR/Cas9 genomic editing approach to deplete METTL3, METTL14, VIRMA and WTAP in Akata (EBV+) cells. Consistent with our prediction, depletion of METTL3, METTL14, VIRMA and WTAP all facilitated the expression of EBV ZTA and RTA (**Figure 8A, 8B & 8D**, lanes 2, 3, 5, 6 vs 8-9; **Figure 8C**, lanes 2, 4, 6 vs 8). Because YTHDF1 and YTHDF3 share redundant role with YTHDF2, we knocked out these two genes using CRISPR/Cas9 genomic editing approaches. Interestingly, depletion of YTHDF3 rather than YTHDF1 promoted EBV ZTA and RTA expression (**Figures 8E and S14E**). In addition, depletion of the major m^6^A eraser ALKBH5 did not affect EBV protein accumulation (**Figure S14F**).

**Figure 8.**
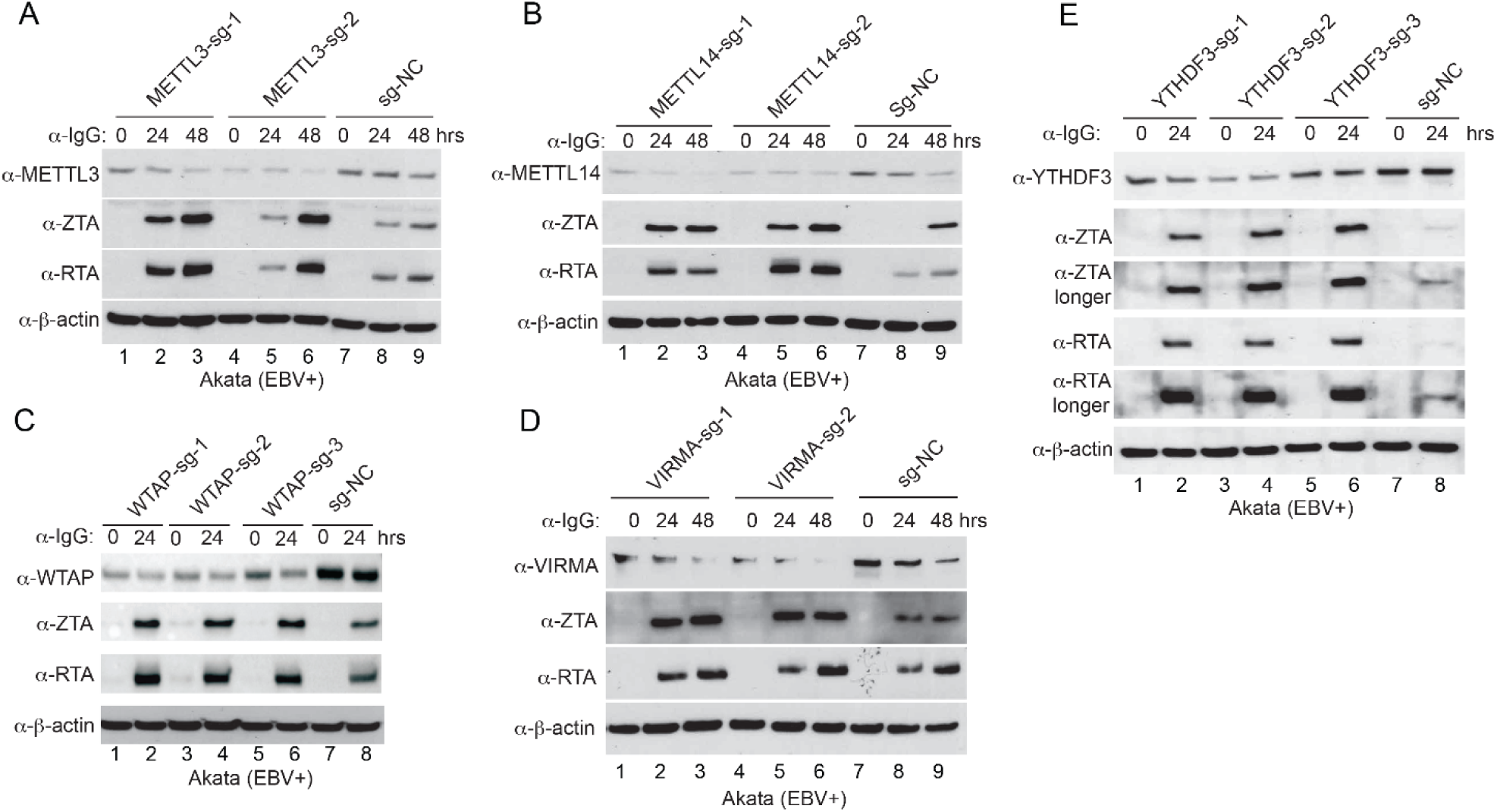
Depletion of m^6^A writers and reader YTHDF3 promotes EBV reactivation. (A-E) Akata (EBV+) cells were used to establish stable cell lines using 2-3 different guide RNA constructs targeting METTL3 (A), METTL14 (B), WTAP (C), VIRMA (D) and YTHDF3 (E) and a non-targeting control (sg-NC). The cells were untreated or lytically induced with anti-IgG-mediated BCR activation. Cellular and viral protein expression levels were monitored by Western Blot using antibodies as indicated. See also Figures S14 and S15.

To further confirm the key roles of m^6^A writers and readers in EBV gene expression and viral life cycle, we measured *ZTA* and *RTA* gene expression by RT-qPCR and virion-associated DNA copy numbers by qPCR for cells with individual gene depleted by CRISPR/Cas9. We found that the depletion of METTL3, METTL14, WTAP, VIRMA and YTHDF3 all promoted viral gene expression (**Figure S15A, S15C, S15E, S15G and S15I**) and facilitated viral replication upon lytic induction (**Figure S15B, S15D, S15F, S15H and S15J**).

All results together suggested that, in addition to m^6^A readers, disruption of the m^6^A writer complex by caspases further fosters EBV reactivation upon lytic induction.

## Discussion

Caspase activation and m^6^A RNA modification pathway have documented essential roles in viral infections. However, it is unknown whether caspases regulate any members of the m^6^A machinery to foster viral infection. Our study established an elegant regulation model about multiple members of the m^6^A RNA modification pathway by caspase mediated cleavage, which play a crucial role in regulating the reactivation of EBV (**Figure 9**).

**Figure 9.**
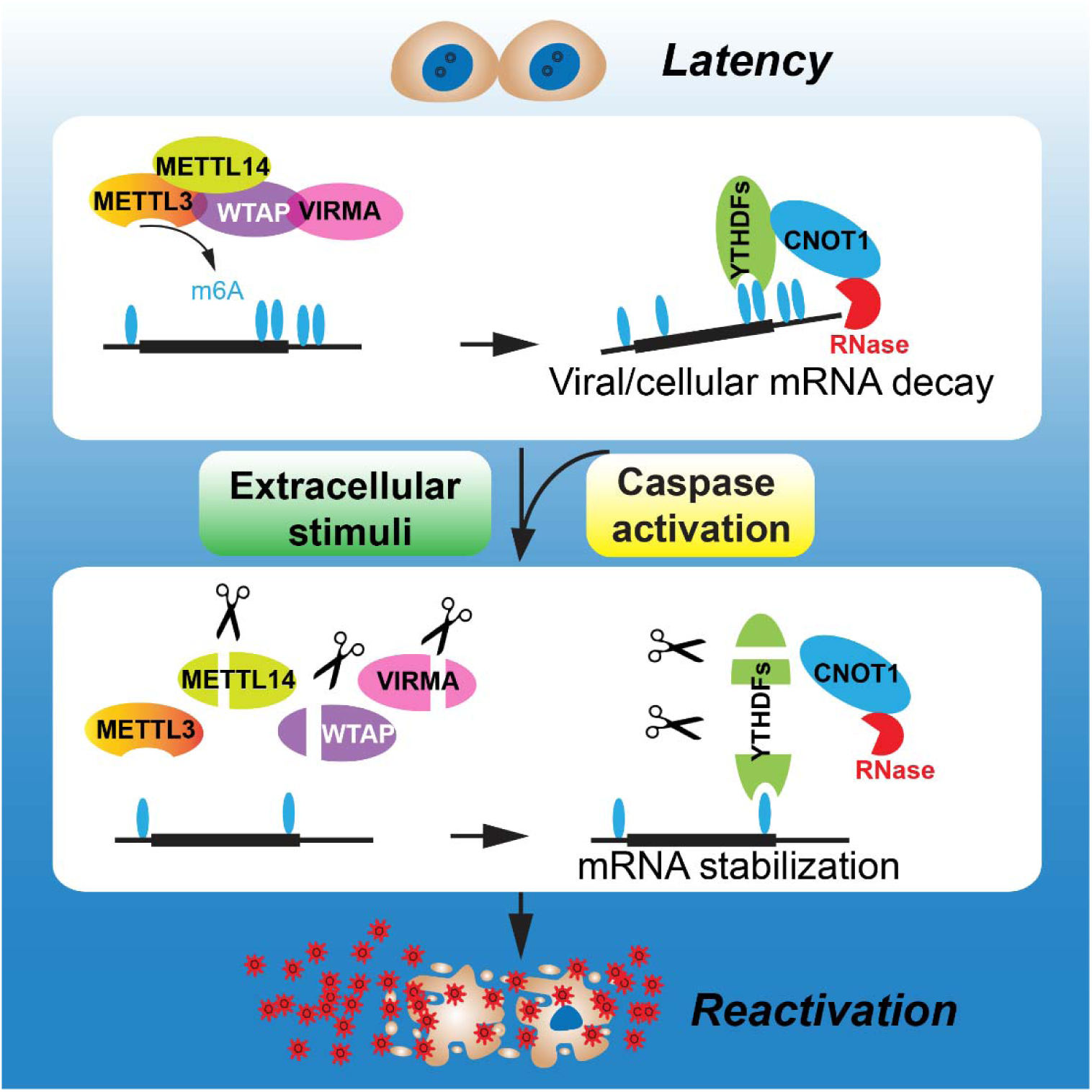
Model: Caspase-mediated cleavage of m^6^A pathway writers and readers facilitates EBV reactivation. During latency, m^6^A writers deposit the methyl group onto key viral and cellular mRNAs which are subsequently destabilized by m^6^A readers. Upon reactivation, cellular caspases are activated. On one hand, caspases cleave the writers to limit the m^6^A modification process, and on the other hand, caspases cleave the readers to limits RNA decay by CNOT-CCR4 complex, which together foster EBV reactivation and drive the production of massive amounts of viruses.

In this study, we first identified 16 putative caspase substrates by searching the human proteome database using two unique cleavage motifs derived from PIAS1 (**Figure 1**). The m^6^A reader YTHDF2 was selected for in-depth analysis considering its critical role in the life cycle of diverse viruses, including EBV and KSHV (**Figure 2**) (25, 28, 29).

Consistent with our prediction, YTHDF2 was indeed cleaved on D367 within the NGVD*G motif shared with PIAS1. Moreover, we discovered a second cleavage site (D166) within the motif AMID*G that is unique for YTHDF2 and not present in any other human proteins (**Figure 3**). Both cleavage sites, especially D166, are evolutionarily conserved across a diverse group of vertebrates ranging from human to zebrafish, highlighting that maintaining these cleavage sites during evolution is important for normal cellular processes (**Figures 3 and S4**).

In addition to YTH domain binding to m^6^A, YTHDF2 contains a low complexity domain with disorder-promoting amino acids in the N-terminal and central parts. Our 3D structure modeling on full-length YTHDF2 clearly shows two cleavage sites on the surface area with flexible turns (**Figure 3J**). Because the low complexity domain is responsible for YTHDF2-mediated phase separation to form liquid droplets with high concentration of protein and RNA (21–23), our predicted protein structure model will provide valuable insights into the regulation of YTHDF family proteins under normal conditions.

The importance of the YTHDF2 cleavage in EBV replication is further supported by the following observations. First, cleavage-resistant YTHDF2 strongly inhibits EBV replication upon lytic induction. Second, overexpression of cleavage-mimicking YTHDF2 fragments promoted the accumulation of EBV immediate-early proteins. Finally, the interaction between YTHDF2 and CNOT1 was diminished upon caspase-mediated cleavage of YTHDF2, limiting the efficient recruitment of CCR4-NOT complex by YTHDF2 (**Figure 5**).

Although YTHDF2 regulation of viral transcripts stability contributes to viral replication (**Figures 2** **& S10**) (25, 28, 29), the extent to which cellular gene regulation by YTHDF2 plays in this process has been largely unexplored. By analyzing YTHDF2 targets transcripts we identified *CASP8* as a putative YTHDF2 target involved in EBV replication (**Figures 6 and S11**). We demonstrated that the *CASP8* transcript is modified by m^6^A and bound by YTHDF2 and that depletion of YTHDF2 promotes *CASP8* mRNA level, caspase-8 protein level, and hence its activation and subsequent cleavage of PIAS1 favoring EBV lytic replication (**Figure 6**).

Intriguingly, we found 15 putative m^6^A sites within the second to last exon. Among them, 5 sites are conserved across a diverse group of species. Although DNA sequences differ significantly across species, our discovery of conserved m^6^A sites in regulating *CASP8* stability highlights the importance of m^6^A RNA modification in controlling the basal levels of key cellular genes during evolution (**Figure 6**). The importance of caspase-8 activation in YTHDF2-depleted cells was further supported by the evidence that caspase-8 inhibition blocks PIAS1 degradation and EBV replication (**Figure S11D-S11G**). Therefore, our discovery of *CASP8* mRNA targeting by YTHDF2 and YTHDF2 cleavage by caspase-8 has established an elegant feedback regulation mechanism controlling the switch of EBV from latent to lytic phase (**Figure 6**).

Importantly, our work on YTHDF2 led to the discovery of multiple m^6^A RNA modification pathway proteins as caspase substrates and EBV restriction factors. The extension of YTHDF2 to YTHDF1 and YTHDF3 as caspase substrates was readily revealed by sequence alignment and cleavage patten analyses (**Figures S12 & S13**).

We provided compelling evidence that METTL14 and WTAP are cleaved by cellular caspases (**Figures 7** **&** **S14**). Interestingly, WTAP is predominately cleaved by caspase-6 on D302 within an evolutionarily conserved motif that differs significantly from canonical cleavage motifs (**Figure S14**). The cleavage pattern of METTL14 suggested that multiple sites are cleaved, possibly including D29 (EASD*S) revealed by a large scale proteomics study (55). It will be interesting to identify all cleavage sites on METTL14 and examine how specific cleavage contributes to the EBV latency and reactivation in the future. Importantly, the discovery of new cleavage motifs (**Figures S13 & S14**) would also guide us to discover novel caspase substrates using the motif searching method (**Figure 1**) and to determine the biological function of protein cleavage in viral life cycle and normal cellular processes.

Our demonstration of m^6^A writers and readers in EBV lytic replication (**Figures 8 and S15**) provided a foundation to further explore how m6A modification of viral and cellular factors contributes to viral life cycle and more broadly normal cellular processes, including cell death and differentiation.

EBV infection of primary human B cells gradually establishes latency by reprograming cellular environment. Considering the role of YTHDF2 in controlling EBV latency, we also extracted the RNA and protein level data from the transcriptomic and proteomic analyses of EBV infection of primary human B cells by the Hammerschmidt group and the Gewurz group (56, 57), respectively. Inherently, we found that EBV infection led to enhanced YTHDF2 RNA and protein expression and reduced CASP8 expression (**Figure S16**). There were also a transient increase and then gradual decrease of EBV latent and lytic gene expression, suggesting that YTHDF2 may also control their expression during latency establishment (**Figure S16**).

There are several limitations in our study. For examples, overexpression of individual fragment may not represent the real cleavage situation and therefore may lead to enhanced phenotypes that we observed. The existence of several fragments in real cleavage situation may coordinate together to promote viral replication. In addition to *CASP8*, the expression of epigenetic regulators, including histone modifiers and transcription factors, may be altered upon the depletion of YTHDF2 and other m^6^A pathway genes (58). Indeed, the transcripts of histone acetyltransferases p300/CBP and histone methyltransferase EZH2 have been shown to be regulated by m^6^A modifications (59, 60). All these factors have been implicated in γ-herpesvirus latency and reactivation (61–63). The manipulation of m^6^A pathway regulators may also change chromatin regulatory RNAs (64). The chromatin regulatory RNAs may also control the viral and cellular chromatin status important for EBV latency and/or reactivation. METTL3 and YTHDC1 were reported to regulate heterochromatin formation via RNA m^6^A methylation in mouse embryonic stem cells (65). The crosstalk between epitranscriptomics and epigenetics in herpesvirus latency and lytic replication is an exciting area to be explored in the future.

In summary, our work has illustrated the value of a motif-based searching approach for the discovery of novel caspase substrates that are critical for viral replication. We have uncovered a unique caspase regulation mechanism for the m^6^A RNA modification pathway, which is essential for the reactivation of a ubiquitous tumor virus. The discovery of conserved caspase cleavage motifs will guide us to discover novel caspase substrates with broad biological significance, provide valuable insights into the regulation of viral life cycle, and illuminate the key molecular mechanisms controlling normal cellular processes and disease progression.

## Materials and Methods

### Cell Lines and Cultures

The Akata (EBV+) and Akata-4E3 (EBV-), P3HR-1 and SNU-719 cells were cultured in RPMI 1640 medium supplemented with 10% FBS (Cat# 26140079, Thermo Fisher Scientific) in 5% CO_2_ at 37°C. 293T cells were cultured in DMEM medium supplemented with 10% FBS (Cat# 26140079, Thermo Fisher Scientific) in 5% CO_2_ at 37 °C. All cell line information is listed in **Table S1**.

### Plasmids Construction

Halo-V5-YTHDF2 (full length and truncation mutants), Halo-V5-METTL13, Halo-V5-METTL14 and Halo-V5-WTAP were cloned into pHTN HaloTag CMV-neo vector (Cat# G7721, Promega) using Gibson assembly methods as previously described (7, 38). Halo-V5-YTHDF2 mutant (D166A/D367A) and Halo-V5-WTAP mutants (D301A and D301A/D302A) were generated using the QuikChange II site-directed Mutagenesis Kit (Stratagene) following the manufacturer’s instructions. All primers are listed in **Table S1**.

### Target Gene Depletion by CRISPR/Cas9 Genome Editing

To deplete YTHDF2, EIF4H, MAGEA10, SORT1, MTA1, EHMT2, METTL3, METTL14, WTAP, VIRMA, YTHDF1, YTHDF3 and ALKBH5, two or three different sgRNAs were designed and cloned into lentiCRISPR v2-Puro vector (a gift from Feng Zhang; Addgene plasmid # 52961) (66) or lentiCRISPR v2-Blast vector (a gift from Mohan Babu, Addgene plasmids #83480) or lentiCRISPR v2-Hygro vector vector (a gift from Joshua Mendell, Addgene plasmids #91977) (67). Packaging 293T cells were transfected with targeted gene sgRNAs or negative control (non-targeting sgRNA-NC) and helper vectors (pMD2.G and psPAX2; gifts from Didier Trono; Addgene plasmid #s 12259 and 12260, received via Yue Sun) using Lipofectamine 2000 reagent (Cat# 11668019, Life Technologies). Medium containing lentiviral particles and 8 mg/mL polybrene (Sigma-Aldrich, St. Louis) was used to infect Akata (EBV+) cells, P3HR-1 cells and SNU-719 cells. The stable cell lines were selected and maintained in RPMI medium supplemented with 2 μg/mL puromycin, 100 μg/mL hygromycin, or 10 μg/mL blasticidin.

To knockout three caspases (CASP3/CASP8/CASP6), first, one sgRNA targeting CASP3 (CASP3-sg1) was cloned into lentiCRISPR v2-Puro vector and Akata (EBV+) cells were used to create a CASP3-depleted cell line, Akata (EBV+)-CASP3-sg1, by CRISPR/Cas9 (puromycin-resistant). Second, one sgRNA targeting CASP8 (CASP8-sg1) and one control sgRNA(sg-NC) were cloned into lentiCRISPR v2-Blast vector. The constructs were packaged, and lentiviral particles were used to infect Akata (EBV+)-CASP3-sg1 and control Akata (EBV+)-sg-NC cells to generate Akata (EBV+)-CASP3-sg1/CASP8-sg1 and Akata (EBV+)-sg-NC/sg-NC cell lines. Third, two different sgRNAs targeting CASP6 (CASP6-sg1 and CASP6-sg2) and one control sgRNA (sg-NC) were designed and cloned into lentiCRISPR v2-Hygro vector. The constructs were packaged, and lentiviral particles were used to infect Akata (EBV+)-CASP3-sg1/CASP8-sg1 and control Akata (EBV+)-sg-NC/sg-NC cell line. The established Akata (EBV+)-CASP3-sg1/CASP8-sg1/CASP6-sg1, Akata (EBV+)-CASP3-sg1/CASP8-sg1/CASP6-sg2, and control Akata (EBV+)-sg-NC/sg-NC/sg-NC were maintained in RPMI medium supplemented with 100 μg/mL hygromycin, 2 μg/mL puromycin and 10 μg/mL blasticidin. The target guide RNA sequences are listed in **Table S1**.

Similarly, to create a *YTHDF2/CASP8*-double knockout cell line, first, one sgRNA targeting *CASP8* (CASP8-sg1) and control sg-RNA in lentiCRISPR v2-Blast vector were used to create *CASP8*-knockout and control cell lines [Akata (EBV+)-CASP8-sg1 and Akata (EBV+)-sg-NC (blasticidin-resistant)]. Second, and lentiviral particles carrying one sgRNA targeting *YTHDF2* (YTHDF2-sg1, puromycin-resistant) were used to infect Akata (EBV+)-CASP8-sg1 and control Akata (EBV+)-sg-NC cell lines. The established Akata (EBV+)-CASP8-sg1/YTHDF2-sg1 and Akata (EBV+)-sg-NC/YTHDF2-sg1 cell lines were maintained in RPMI medium supplemented with 2 μg/mL puromycin and 10 μg/mL blasticidin. The target guide RNA sequences are listed in **Table S1**.

### Lentiviral Transduction of YTHDF2

The pLenti-C-Myc-DDK-P2A-BSD and pCMV6-Entry-YTHDF2 were purchased from Origene. The specific variants were generated by site-directed mutagenesis in pCMV6-Entry-YTHDF2 using the QuikChange II site-Directed Mutagenesis Kit (Cat# 200521, Stratagene) according to the manufacturer’s instructions. The truncated YTHDF2 were cloned into pCMV6-Entry vector using Gibson assembly methods as we previous described (7, 38). Subsequently, the WT, mutated and truncated YTHDF2 in pCMV6-Entry vector were digested using AsiSI and MluI and subcloned into the pLenti-C-Myc-DDK-P2A-BSD vector. To prepare lentiviruses, 293T cells were transfected with lentiviral vector containing the target gene and the help vectors (pMD2.G and psPAX2) using Lipofectamine 2000 reagent. The supernatants were collected 48 hs post-transfection to infect Akata (EBV+) cells or SNU-719 cells and then stable cell lines were selected in RPMI medium containing 10 μg/mL blasticidin.

For YTHDF2 reconstitution, we first transduced the Akata (EBV+) cells with lentiviruses carrying vector control, WT and mutant (D166A/D367A) YTHDF2 to establish cell lines in RPMI medium containing 10 μg/mL blasticidin. We then depleted endogenous YTHDF2 by lentiviral transduction of YTHDF2-sg2 into these three cell lines. The obtained stable cell lines were selected in RPMI medium supplemented with 2 μg/mL puromycin and 10 μg/mL blasticidin.

### Lentiviral Transduction of CASP8

The full-length *CASP8* were cloned into pCMV6-Entry vector using Gibson assembly methods as we previous described (7, 38). Subsequently, the full-length CASP8 in pCMV6-Entry vector were digested using AsiSI and MluI and subcloned into the pLenti-C-Myc-DDK-P2A-BSD vector. To prepare lentiviruses, 293T cells were transfected with lentiviral vector containing *CASP8* gene and the help vectors (pMD2.G and psPAX2) using Lipofectamine 2000 reagent. The supernatants were collected 48 hs post-transfection to infect Akata (EBV+) cells and then Akata (EBV+)-*CASP8* stable cell line was selected in medium containing 10 μg/mL blasticidin.

### Lytic Induction and Cell Treatment

For lytic induction in Akata (EBV+) cell lines, the cells were treated with IgG (1:200, Cat# 55087, MP Biomedicals) for 0 to 48 hrs. Akata-4E3(EBV-) cells were treated similarly as controls. To induce the EBV lytic cycle in P3HR-1 cells, the cells were triggered with TPA (20 ng/ml) and sodium butyrate (3 mM) for 0 to 48 hrs. For EBV lytic induction in SNU-719 cells, the cells were treated with gemcitabine (1 μg/mL) or TPA (20 ng/ml) and sodium butyrate (3 mM) for 0 to 48 hrs. For caspase inhibition assay, Akata (EBV+), P3HR-1 and SNU719 cells were untreated or pretreated with caspase inhibitors (50 μM) for 1 hr and then treated with lytic inducers for additional 24 or 48 hrs. For apoptotic induction, Akata (EBV+), P3HR-1 and SNU719 cells were treated with Taxol at different concentrations for 24 or 48 hrs. All key reagent information is listed in **Table S1**.

### Cell Lysis and Immunoblotting

Cell lysates were prepared in lysis buffer supplemented with protease inhibitors (Roche) as described previously. Protein concentration was determined by Bradford assay (Biorad). The proteins were separated on 4-20% TGX gels (Biorad) and transferred to PVDF membranes using semi-dry transfer system. Membranes were blocked in 5% milk and probed with primary and horseradish peroxidase-conjugated secondary antibodies. All antibody information is listed in **Table S1**.

### Protein Expression and Purification

Halo-V5-YTHDF2 (WT and D166/D367A mutant), Halo-V5-METTL3, Halo-V5-METTL14 and Halo-V5-WTAP (WT, D301A and D301/D302A mutants) proteins were expressed and purified as previously described (68). Briefly, Halo-tagged plasmids were transfected into 293T cells at 50-60% confluence. Two T175 flasks of transfected cells were harvested 48 hrs post-transfection at 100% confluence and lysed with 25 ml HaloTag Protein Purification Buffer (50 mM HEPES pH7.5, 150 mM NaCl, 1mM DTT, 1mM EDTA and 0.005% NP40/IGEPAL CA-630) with Protease Inhibitor Cocktail. Halo-V5-tagged proteins were enriched using the Halo-tag resin and proteins were eluted from the resin by washing 3 times with 0.5 ml HaloTag Protein Purification Buffer containing 20 μl Halo-TEV protease. The eluted proteins were stored at -80 °C for further use.

### Reverse transcription and quantitative PCR (RT-qPCR)

Total RNA was isolated from cells with Isolate II RNA Mini Kit (Bioline) according to the manufacturer’s instructions. Total RNA was reverse transcribed into cDNA using the High Capacity cDNA Reverse Transcription Kit (Invitrogen). qPCR was performed using Brilliant SYBR Green qPCR master mix (Agilent Technology) with specific primers listed in **Table S1**. The relative expression of target mRNA was normalized by β*-actin* or *HPRT1* expression level.

### EBV DNA Detection

To measure EBV replication, cell associated viral DNA and virion-associated DNA were determined by qPCR analysis. For intracellular viral DNA, total genomic DNA was extracted using the Genomic DNA Purification Kit (Cat# A1120, Promega). For extracellular viral DNA, the supernatant was treated with RQ1 DNase at 37 °C for 1 hr, deactivation by RQ1 DNase. stop solution, followed by releasing virion associated DNA with proteinase and SDS treatment as previously described (6). The DNA was then purified by phenol/chloroform/isoamyl alcohol extraction. The relative viral DNA copy numbers were determined by qPCR using primers to *BALF5* gene. The reference gene β*-actin* was used for data normalization.

### *In vitro* Caspase Cleavage Assay

Purified V5-YTHDF2 (WT and mutants), V5-WTAP (WT and mutants), WT V5-METTL3 and WT V5-METTL14 were incubated with individual caspase in caspase assay buffer (50 mM HEPES, pH7.2, 50 mM NaCl, 0.1% Chaps, 10 mM EDTA, 5% Glycerol and 10mM DTT) at 37 °C for 2 hrs with gentle agitation. Reactions were stopped by boiling in 2x SDS sample buffer and samples were analyzed by Western blot.

### Immunofluorescence Assay

Akata EBV (+) cells were grown and treated with IgG (1:200) for 24 and 48 hrs in 6-well culture dish. Cell suspensions were collected and centrifuged. Cell pellets were then re-suspended with PBS (Phosphate-buffered saline, pH 7.4) and transferred to a 12-well culture dish containing poly-L-Lysine coated coverslips. SNU-719 cells were grown in sterilized coverslips in a 12 well culture dish and then treated with TPA (20 ng/ml) and sodium butyrate (3 mM) for 24 and 48 hrs. The cells were fixed with 4% paraformaldehyde and washed 3 times with PBS. Cells were stained with propidium iodine and then washed 5 times with PBS, followed by permeabilization with 0.25% Triton X-100 for 5 mins and blocking with 3% bovine serum albumin (BSA) for 1 hr at room temperature. The cells were then incubated with rabbit anti-YTHDF2 antibody (1:250, Proteintech) in 3% BSA for 2 hrs at room temperature. The cells were washed 3 times with PBS and incubated with Alexa Fluor 488-conjugated anti-rabbit secondary antibody (1:500) for 1 hrs at room temperature. For co-immunostaining, the cells were fixed with paraformaldehyde and permeabilized with 0.25% Triton X-100. The cells were blocked with 3% BSA and incubated with mouse anti-EBV-EAD (1:250, Millipore) and rabbit anti-YTHDF2 (1:250, Proteintech) antibodies in 3% BSA for 2 hrs at room temperature. The Cells were washed 3 times with PBS and incubated with Alexa Fluor 488-conjugated anti-mouse and Alexa Fluor 568-conjugated anti-rabbit secondary antibodies (1:500) for 1 hr at room temperature. The nuclear DNA was stained using DAPI (1μg/mL, Thermo Fisher Scientific). The coverslips were mounted on slides with antifade reagent, and the immunofluorescence was observed using Axio Observer 7 fluorescence microscope (Carl Zeiss). All key reagent information is listed in **Table S1**.

### Immunoprecipitation (IP) Assay

293T cells (50-60% confluence) were co-transfected with Halo-V5 tagged WT or truncated YTHDF2 and HA tagged CNOT1-SH domain using Lipofectamine 2000. The cells were harvested 48 hrs post-transfection and lysed in RIPA lysis buffer (50 mM Tris-HCl, 150 mM NaCl, 1% NP40, 1% deoxycholate, 0.1% SDS and 1 mM EDTA) containing protease inhibitor cocktail (Cat# 11836153001, Roche) and phosphatase inhibitors (1 mM Na_3_VO_4_ and 1 mM NaF). The IP was carried out as previously described (7, 38).

### RNA-binding Protein Immunoprecipitation

RNA-binding protein immunoprecipitation (RIP) was performed using a Magna RIP kit (Cat# 17-700, Millipore) according to the manufacturer’s protocol. Briefly, Akata (EBV+) cells were either untreated or treated by IgG cross-linking for 24 hrs and then lysed with RIP lysis buffer from the kit. A part of the lysate (10%) was saved as input. The beads were washed with RIP wash buffer, followed by incubation with YTHDF2 antibody (Proteintech, 24744-1-AP) for 30 mins at room temperature and then washed twice with RIP wash buffer. The cell lysate was incubated with antibody-coated beads overnight at 4 °C. The next day, the beads were collected and washed six times with RIP wash buffer. The enriched RNA-protein complex was treated with proteinase K and the released RNA was purified using phenol-chloroform extraction and reverse transcribed for further qPCR analysis using primers listed in **Table S1**.

### M^6^A-RNA Immunoprecipitation (m^6^A RIP) Assay

Total RNA was extracted from Akata (EBV+) cells (untreated or treated by IgG cross-linking for 24 hrs) by TRIzol (Cat# 1596026, Thermo Scientific) and then treated with RQ1 DNase (Cat# M6101, Promega) at 37 °C for 30 mins, followed by RNA re-extraction using TRIzol. 40 μL protein A/G magnetic beads (Thermo Scientific, 88802) were blocked in 1% BSA solution for 2 h, followed by incubation with 10 μg of anti-m^6^A antibody (Millipore, ABE572) at 4 °C for 1 h with rotation. 300 μg purified RNA was added to the antibody-bound beads in IP buffer containing 10 mM Tris-HCL at pH 7.4, 50mM NaCl and 0.1% NP-40 supplemented RNasin plus RNase inhibitor (Cat# N2611, Promega) and incubated overnight at 4 °C with rotation. The same RNA aliquot was used as input. The beads were washed three times with IP buffer, and RNA was eluted twice with IP buffer containing 6.67 mM m^6^A salt (Cat# M2780, Sigma-Aldrich) (200 μL for each elution). The elutes were pooled together and then purified with phenol-chloroform extraction. The immunoprecipitated RNA was reverse transcribed to cDNA for qPCR analysis. The primers were listed in **Table S1**.

### Reporter Cloning and Luciferase Assay

To generate *CASP8*-Exon-7 reporters, we used a psiCheck2 m^6^A-null vector, in which all putative m^6^A sites in the *Renillia* and *Firefly luciferase* genes were mutated (54). The WT *CASP8*-Exon-7, m^6^A Mut1 (5 conserved m^6^A sites were mutated to T) and Mut2 (all putative m^6^A sites were mutate to T) DNA fragments were synthesized using gBlocks (IDT) and were cloned into psiCheck2 m^6^A null vector by Gibson assembly methods using primers and templates listed in **Table S1**.

For luciferase assay, SNU-719 cells and SNU-719 YTHDF2-KD cells were seeded in 12-well plates prior to transfection. The cells were transfected with *CASP8*-Exon-7 luciferase reporters using Lipofectamine 2000 reagent. Forty-eight hours post-transfection, cell extracts were harvested and measured using the Dual-Luciferase assay kit (Promega). Each condition was performed in triplicate.

### Bioinformatic Analysis

The ’multiz100way’ alignment prepared from 100 vertebrate genomes was downloaded from UCSC Genome Browser (http://hgdownload.cse.ucsc.edu/goldenpath/hg38/multiz100way/). The nucleotide sequences around motifs of YTHDF1, YTHDF2, YTHDF3, and WTAP were extracted from ’multiz100way’ alignment by maf_parse of the Phast package (69). The corresponding protein amino acid sequences of each motif were inferred based on the nucleic acid sequence alignments. Sequences containing frameshifting indels were discarded for amino acid alignment to avoid ambiguity. Motif logos were then generated by the WebLogo3 (70).

To analyze the conservation of *CASP8-*Exon-7, we downloaded the phastCons 100-way conservation scores from 100 vertebrate genomes aligned to the human genome (http://hgdownload.cse.ucsc.edu/goldenpath/hg19/phastCons100way/). Given the m^6^A motif (DRACH), we obtained 15 putative m^6^A motif sequences within human *CASP8*-Exon 7 while extracting corresponding sequences from 100 species. After removing sequences containing deletions within the m^6^A motif, we generated 15 motif logos by using the R package ggseqlogo (71).

### Quantification and Statistical Analysis

Statistical analyses employed a two-tailed Student’s t test using Microsoft Excel software for comparison of two groups. A p value less than 0.05 was considered statistically significant. Values were given as the mean ± standard deviation (SD) of biological or technical replicate experiments as discussed in figure legends.

## Acknowledgements

We thank S. Diane Hayward (Johns Hopkins) for providing reagent and cells lines. We thank Stacy Horner (Duke University) for providing a dual luciferase reporter plasmid. We thank Feng Zhang (MIT/Broad), Mohan Babu (University of Regina) and Joshua Mendell (University of Texas Southwestern Medical Center) for sharing the lentiCRISPR v2 plasmids. We also thank Didier Trono (EPFL) for providing the pMD2.G and psPAX2 plasmids.

## Author Contributions

Conceptualization, R.L. and K.Z.; Methodology, K.Z., R.L., Y.M., Y.Z. and J.W.; Investigation, K.Z., R.L., Y.M., F.G.S, Y.Z. and J.W,; Writing – Original Draft, R.L. and K.Z.; Writing – Review & Editing, K.Z., R.L., Y.Z. and J.W.; Funding Acquisition, R.L and J.W.; Visualization, K.Z., R.L., Y.M., Y.Z. and J.W.; Supervision, R.L., and J.W.

## Data and materials availability

All the data needed to evaluate the conclusions in this paper are present in the paper and/or the Supplementary Materials. Additional data related to this paper may be requested from the authors.

---

**Figure S1.**
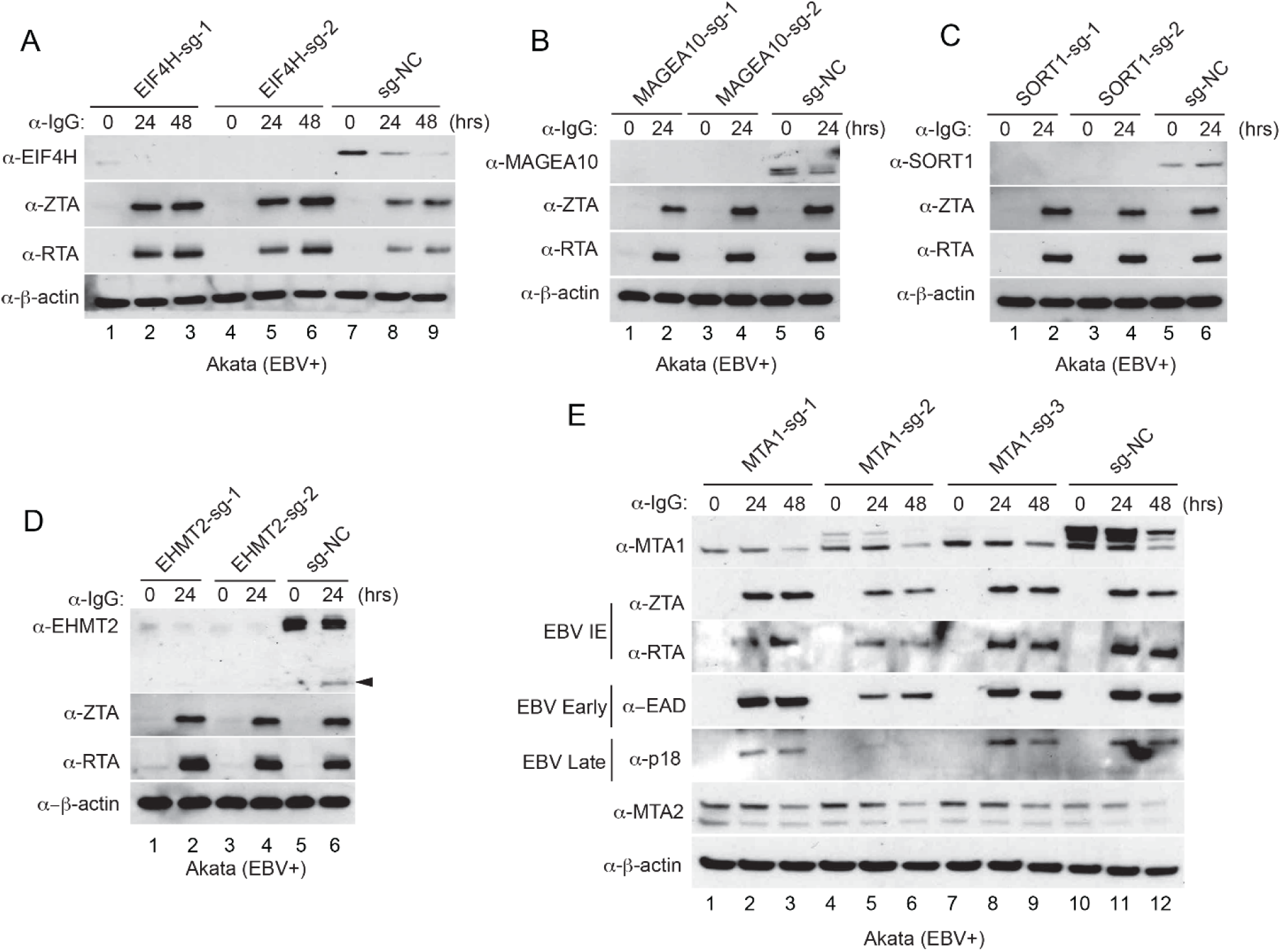
The role of EIF4H, MAGEA10, SORT1, EHMT2, and MTA1 during EBV lytic induction. See also Figure 2. (A-E) Akata (EBV+) cells were used to establish stable cell lines using 2 or 3 different sgRNA constructs and a non-targeting control (sg-NC). The cells were untreated or lytically induced with anti-IgG treatment for 24 or 48 hrs as indicated. Cellular and viral protein expression levels were monitored by Western Blot using antibodies as indicated. (A) EIF4H depletion promotes the expression of EBV ZTA and RTA. (B) MAGEA10 depletion does not affect EBV protein expression. (C) SORT1 depletion does not significantly affect EBV protein expression. (D) EHMT2 depletion does not affect EBV protein expression. Arrowhead denotes cleaved fragments. (E) MTA1 depletion does not uniformly affect EBV protein expression but slightly enhances the expression of its homolog MTA2.

**Figure S2.**
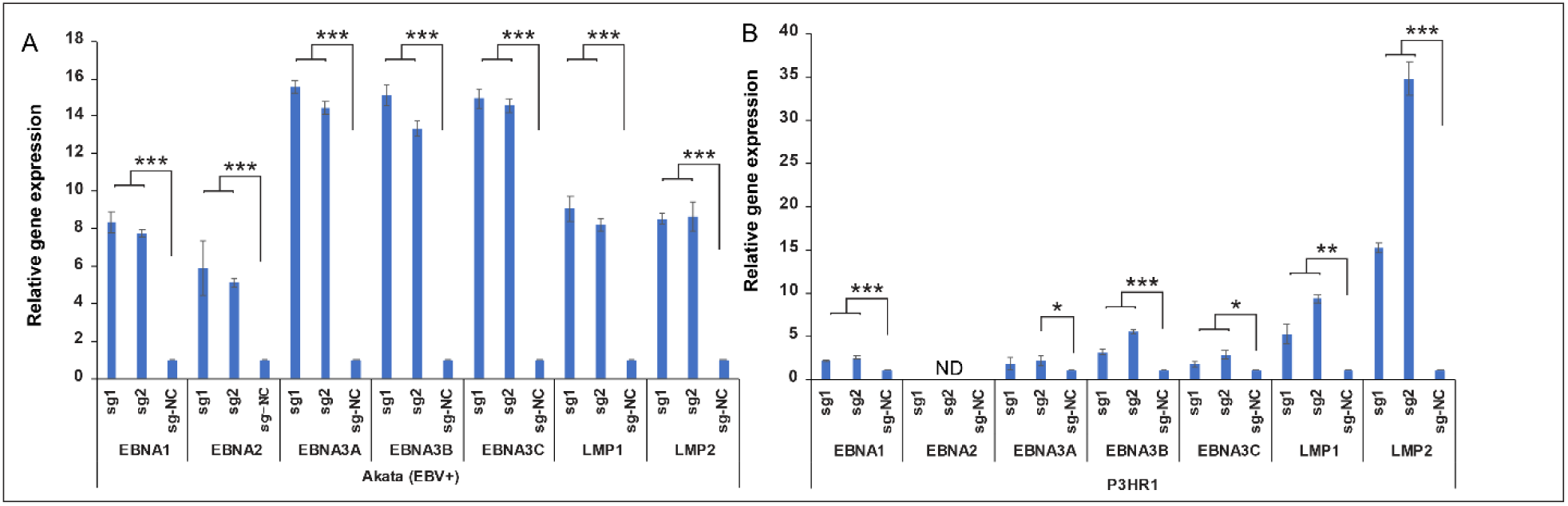
YTHDF2 depletion promotes EBV latent gene expression. See also Figure 2. (**A**) RNAs from YTHDF2-depleted and control Akata (EBV+) cells were extracted and EBV latent genes were analyzed by RT-qPCR. The values of control were set as 1. (B) RNAs from YTHDF2-depleted and control P3HR-1 cells were extracted and EBV latent genes were analyzed by RT-qPCR. The values of control were set as 1. EBV strain in P3HR-1 cells has a deletion that disrupts EBNA2 expression. Error bars indicate ±SD. ND, Not detected. *, p< 0.05; **, p< 0.01; ***, p< 0.001.

**Figure S3.**
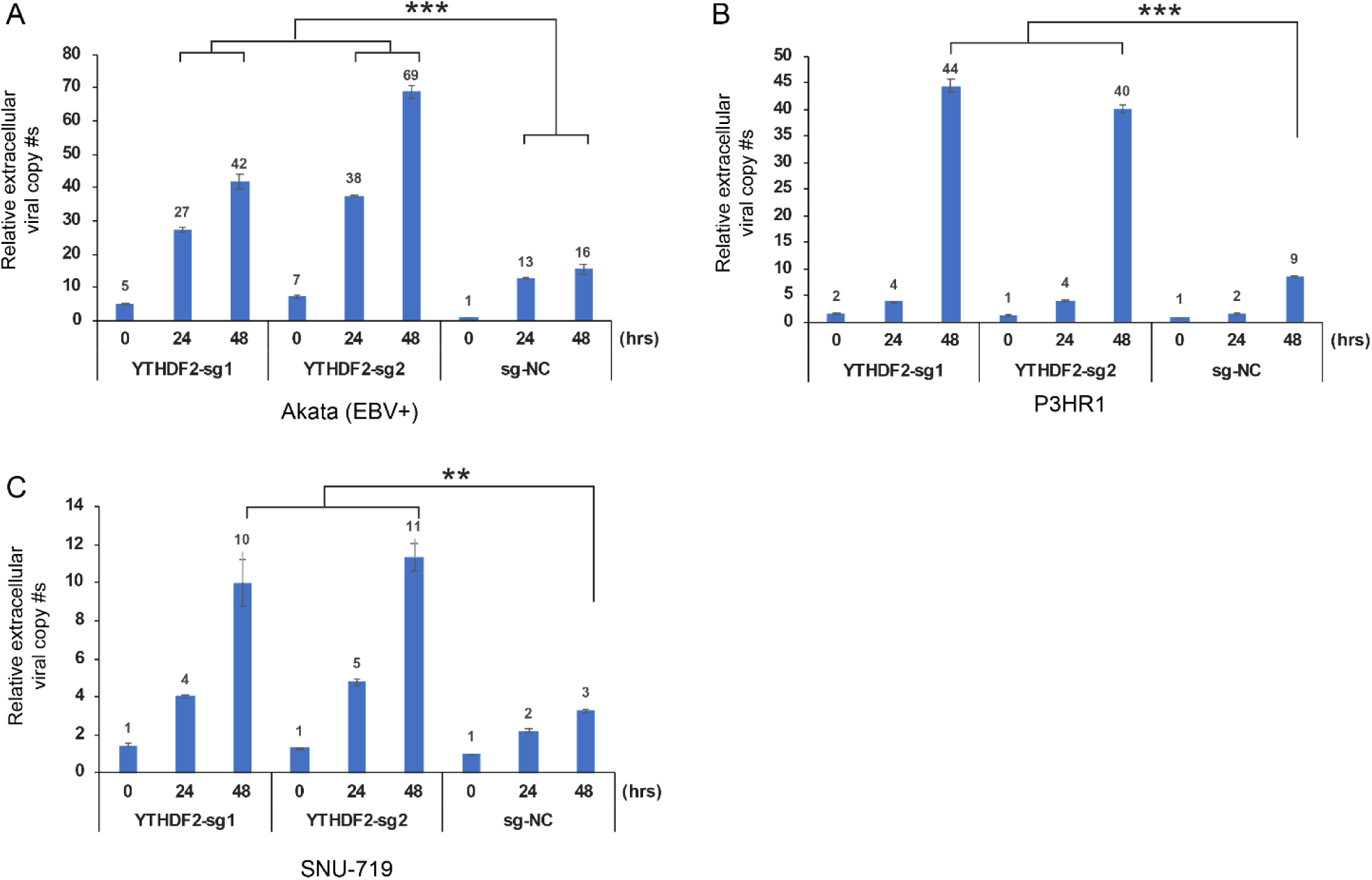
YTHDF2 depletion promotes EBV lytic replication. See also Figure 2. (A) YTHDF2-depleted and control Akata (EBV+) cells were lytically induced with anti-IgG for 0 to 48 hrs. (B) YTHDF2-depleted and control P3HR1 cells were lytically induced with TPA and NaBu for 0 to 48 hrs. (C) YTHDF2-depleted and control SNU-719 cells were lytically induced with TPA and NaBu for 0 to 48 hrs. Extracellular virion DNA from the medium were extracted and then analyzed by qPCR using primers specific to BALF5. The value of vector control at 0 hr was set as 1. Results from three biological replicates are presented. Error bars indicate ±SD. **, p< 0.01; ***, p< 0.001.

**Figure S4.**
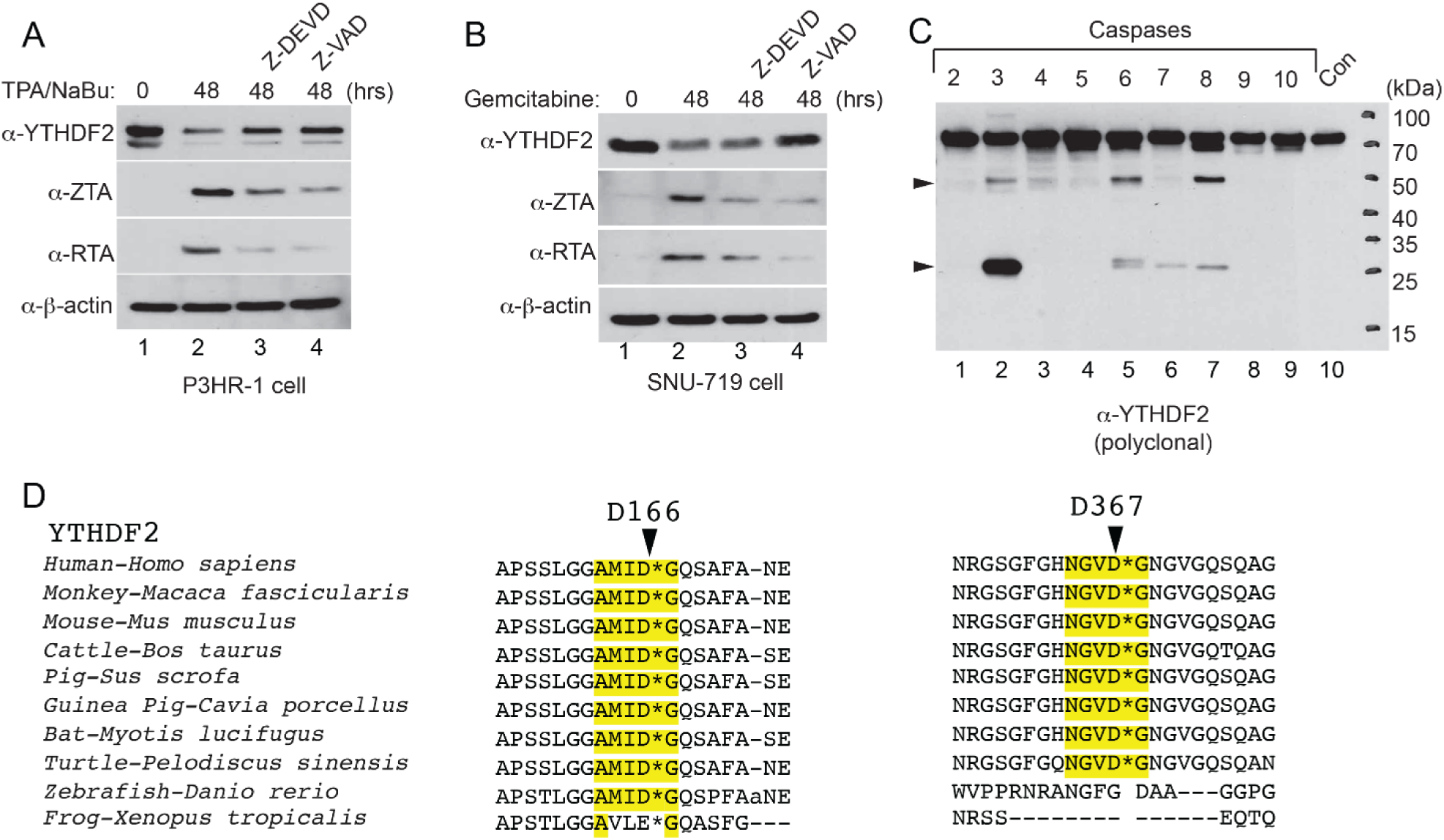
YTHDF2 is cleaved by caspases. See also Figure 3 and Table S2. (A) Caspase inhibition blocks YTHDF2 degradation and viral protein expression. P3HR1 cells were either untreated or pretreated with caspase-3 (Z-DEVD-FMK) or pan-caspase inhibitors (Z-VAD-FMK) (50 μM) for 1 hr and then lytically induced by TPA and sodium butyrate (NaBu) for 48 hrs. YTHDF2 and EBV ZTA/RTA protein levels were monitored by Western Blot as indicated. (B) Caspase inhibition blocks YTHDF2 degradation in SNU-719 cells. SNU-719 cells were either untreated or pretreated with caspase-3 or pan-caspase inhibitors (50 μM) for 1 hr and then lytically induced by gemcitabine (1 μg/ml) for 48 hrs. YTHDF2 and EBV ZTA/RTA protein levels were monitored by antibodies as indicated. (C) Recombinant wild-type YTHDF2 was incubated with individual caspases for 2 hr at 37°C. Western Blot analysis showing YTHDF2 cleavage by caspase-3, -6, -8 and to a lesser extent, caspase-7. Arrowheads denote cleavage fragments. (D) Sequence alignment of YTHDF2 sequences from 10 representative species using the Constraint-based Multiple Alignment Tool (COBALT). The cleavage motifs were highlighted by yellow color.

**Figure S5.**
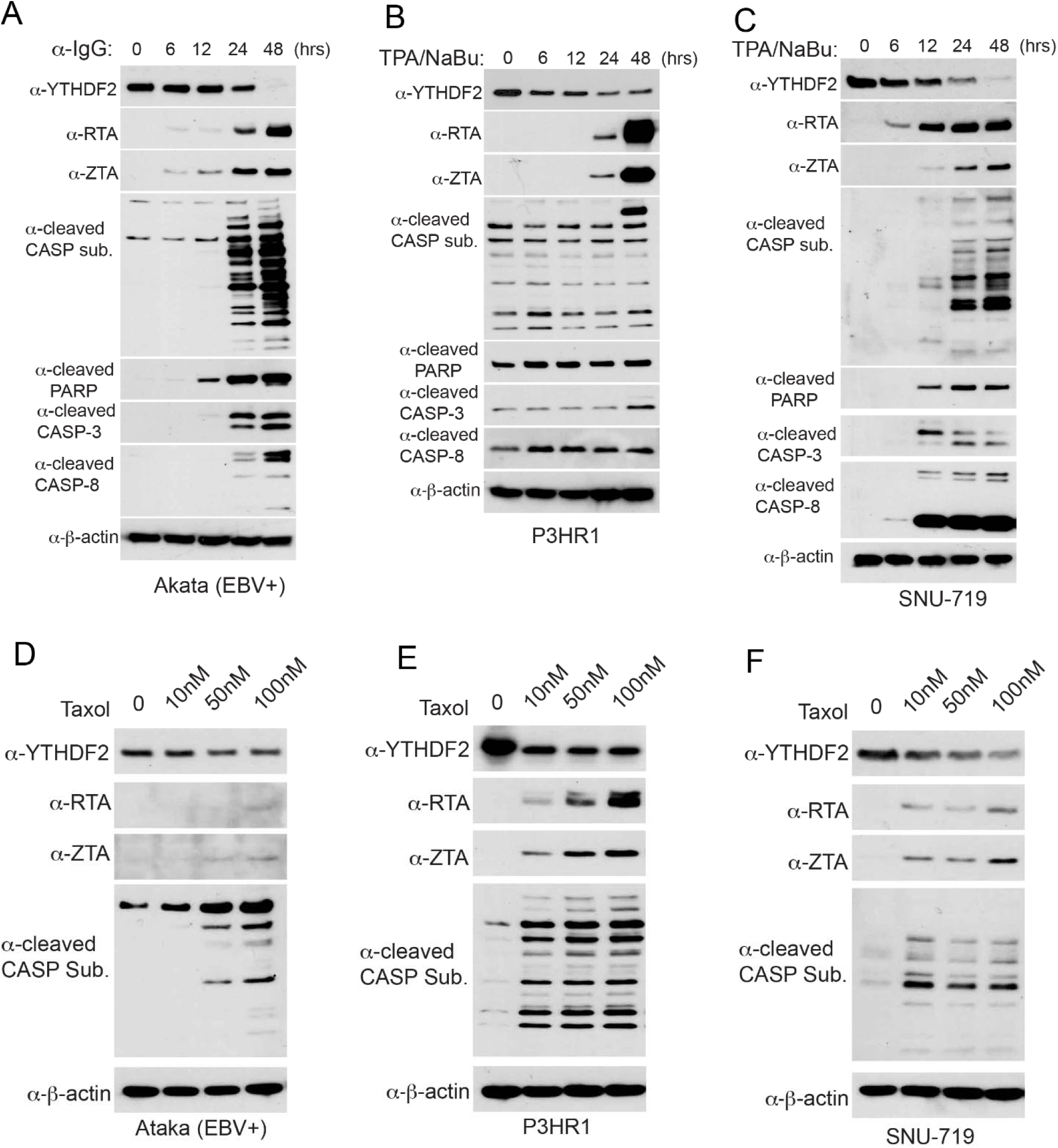
Apoptotic induction promotes YTHDF2 degradation and EBV reactivation. See also Figure 3. (A-C) Akata (EBV+) cells were lytically induced with anti-IgG for 0, 6, 12, 24 and 48 hrs (A). P3HR1 (B) and SNU-719 (C) cells were lytically induced with TPA and NaBu for 0 6, 12, 24 and 48 hrs. YTHDF2, EBV ZTA and RTA, cleaved caspase substrate (CASP sub.), cleaved PARP, cleaved CASP3 and cleaved CASP8 were monitored by Western Blot using antibodies as indicated. β-actin blots were included for loading controls. (D-F) Apoptotic induction by an intrinsic trigger promotes EBV reactivation. Akata (EBV+) (D), P3HR1 (E) and SNU-719 (F) cells were untreated or treated with increasing amount of Taxol for 48 hrs. YTHDF2, EBV ZTA and RTA, and cleaved caspase substrate (CASP sub.) were monitored by Western Blot using antibodies as indicated. β-actin blots were included for loading controls.

**Figure S6.**
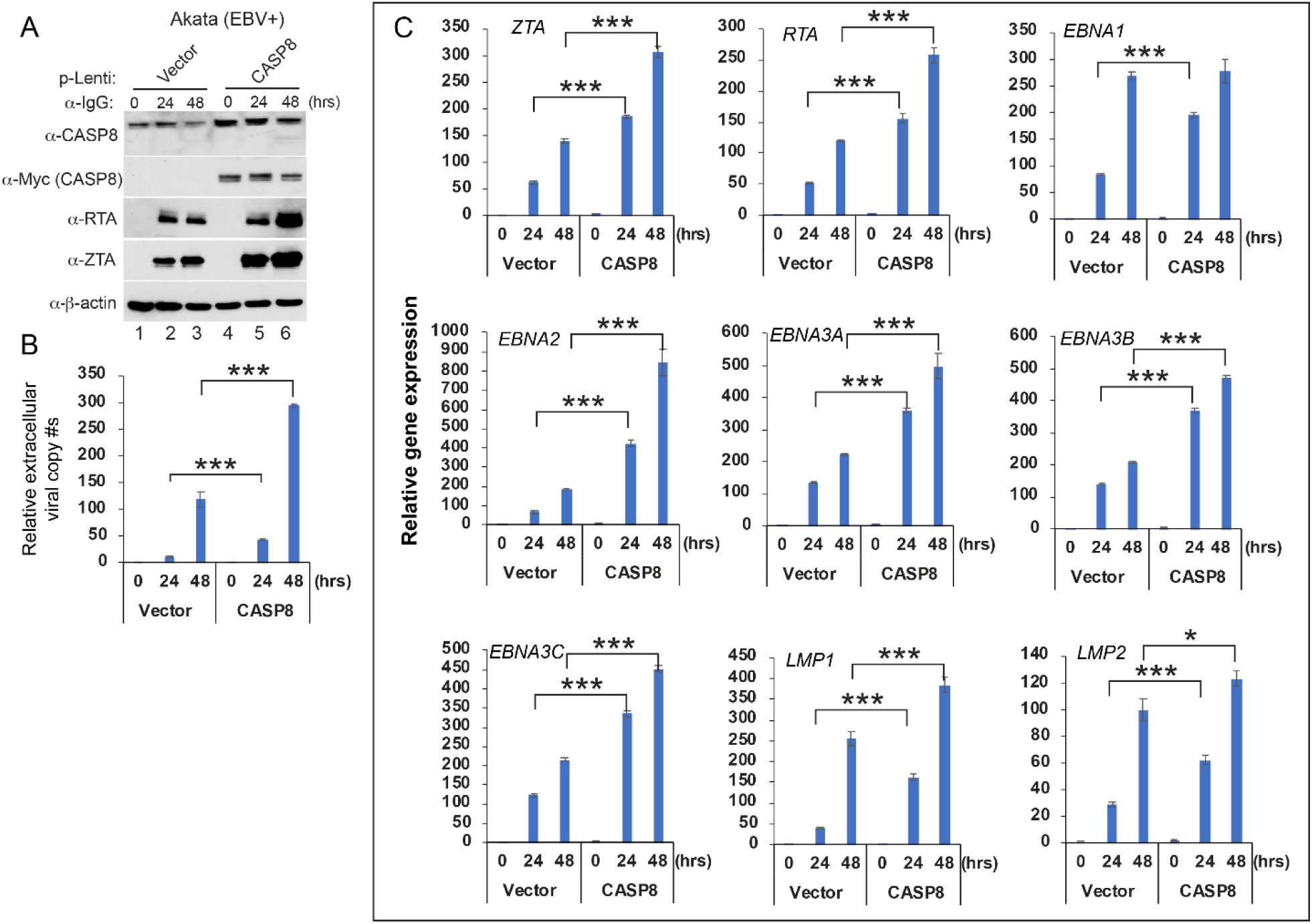
CASP8 overexpression promotes EBV reactivation. See also Figures 3-4. Akata EBV(+) cells were transduced with lenti-vector control or Myc-CASP8 to establish stable cell lines. The cells were treated with anti-IgG for 0, 24 and 48 hrs. (A) CASP8, EBV ZTA and RTA were monitored by Western Blot using antibodies as indicated. β-actin blots were included for loading controls. (B) Extracellular virion DNA from the medium were extracted and then analyzed by qPCR using primers specific to BALF5. The value of vector control at 0 hr was set as 1. (C) Total RNA was extracted and then EBV lytic (ZTA and RTA) and latent (EBNA1, EBNA2, EBNA3A, EBNA3B, EBNA3C, LMP1 and LMP2) genes were analyzed by RT-qPCR. The value of vector control at 0 hr was set as 1 Results from three biological replicates are presented. Error bars indicate ±SD. *, p< 0.05; **, p< 0.01; ***, p< 0.001.

**Figure S7.**
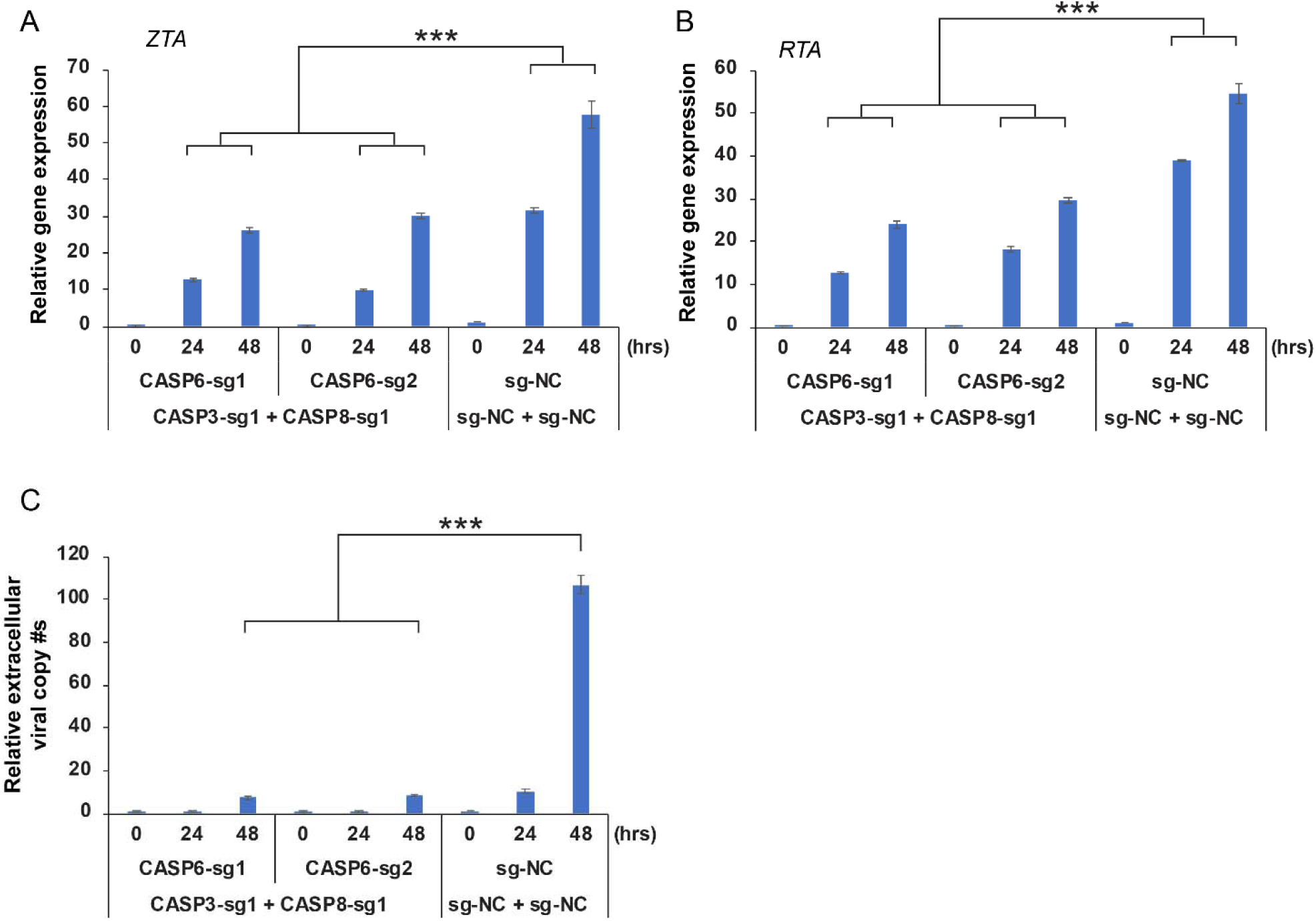
Caspase-3/-6/-8 triple knockout suppresses EBV lytic replication. See also Figure 3. The CASP3/CASP8/CASP6-triply-depleted Akata (EBV+) cells were lytically induced by anti-IgG treatment. (A-B) Total RNA was extracted and then EBV ZTA and RTA mRNA levels were analyzed by RT-qPCR. (C) Extracellular virion DNA from the medium were extracted and then analyzed by qPCR using primers specific to BALF5. The value of control at 0 hr was set as 1. Results from three biological replicates are presented. Error bars indicate ±SD. ***, p< 0.001.

**Figure S8.**
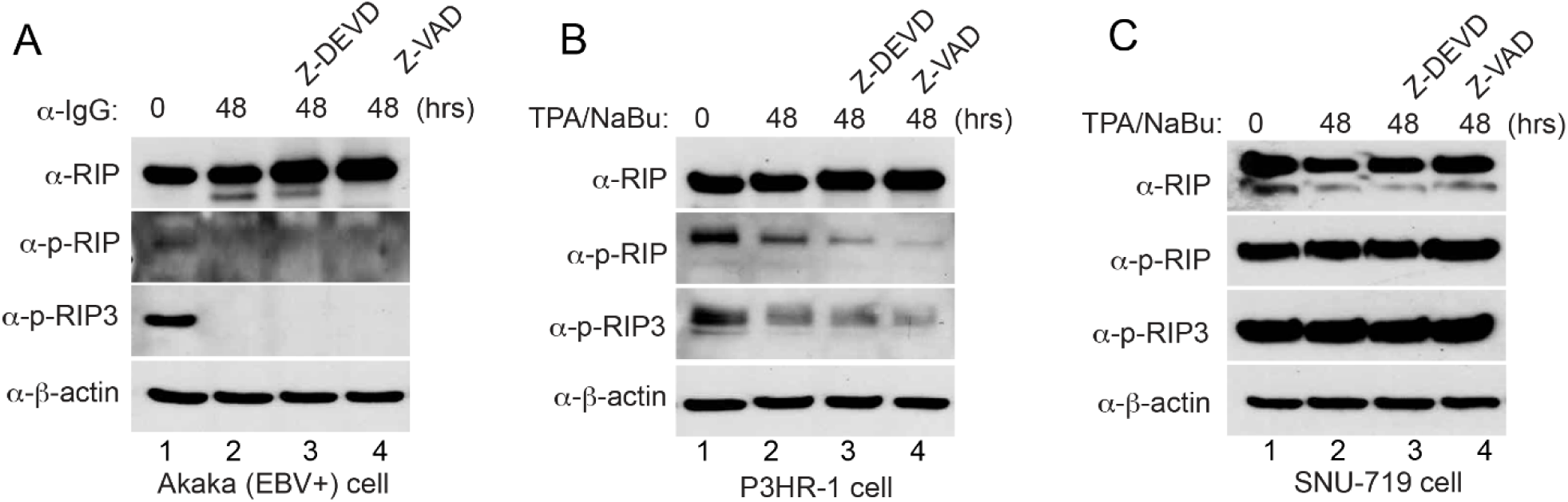
Caspase inhibition does not affect necroptotic cell death pathway. See also Figure 3. The Akata (EBV+) (A), P3HR-1 (B) and SNU-719 (C) cells were either untreated or pretreated with a caspase-3/-7 inhibitor (Z-DEVD-FMK, 50 μM) or pan-caspase inhibitor (Z-VAD-FMK, 50 μM) for 1 hr, and then lytically induced with anti-IgG antibody or TPA/NaBu as indicated for 48 hrs. Western Blot showing the protein levels of RIP, phospho-RIP (p-RIP) and phospho-RIP3 (p-RIP3) using antibodies as indicated. β-actin blots were included as controls.

**Figure S9.**
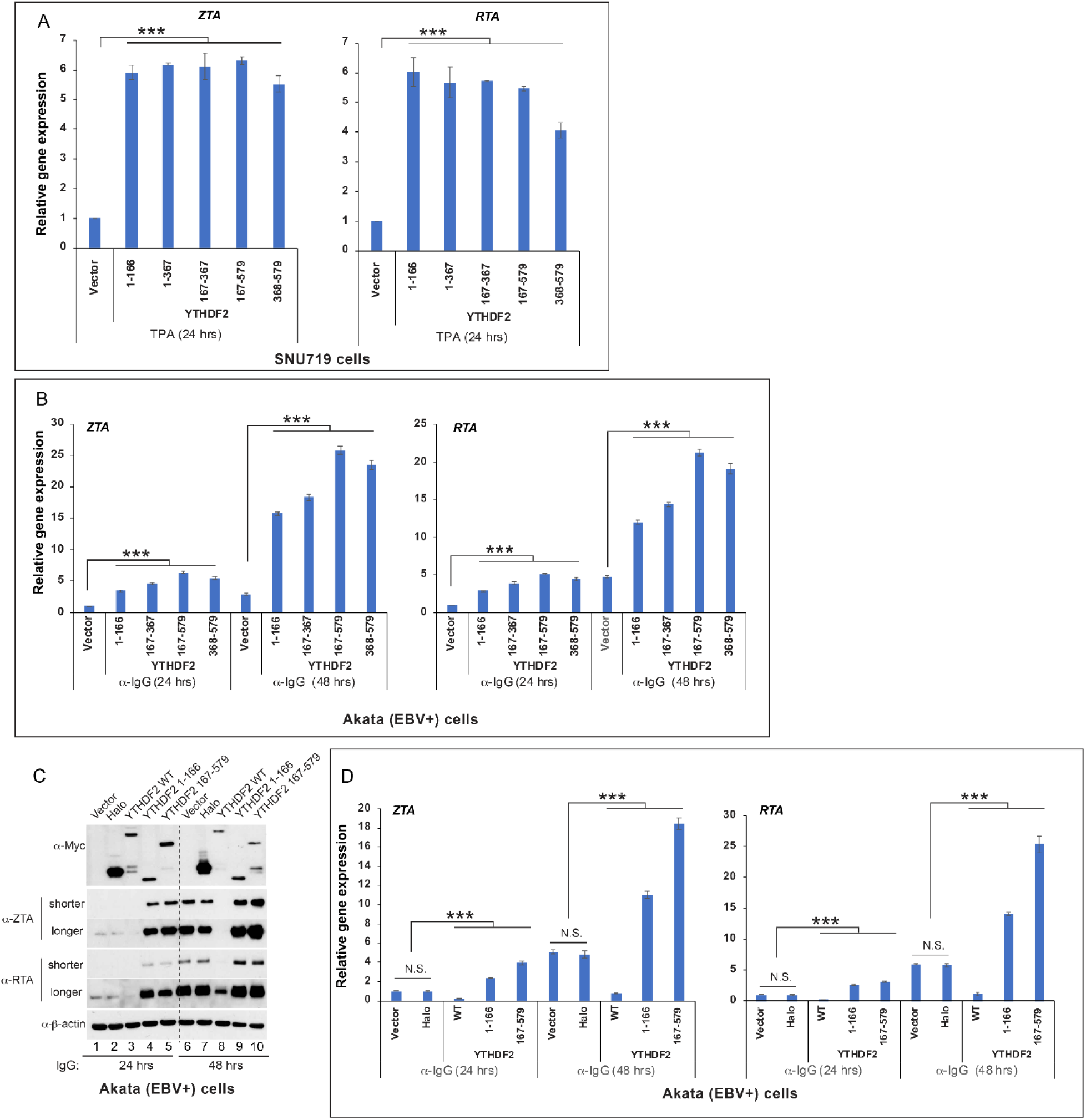
YTHDF2 cleavage fragments promotes EBV lytic gene expression. See also Figure 5. (A) SNU-719 cells were transduced with lentiviruses carrying vector control or individual YTHDF2 fragment to establish stable cell lines (see Figure 5E). RT-qPCR analysis showing EBV ZTA and RTA mRNA levels in these cell lines upon lytic induction by adding TPA (20 ng/ml) for 24 hrs. The value of vector control was set as 1. (B) Akata (EBV+) cells were transduced with lentiviruses carrying vector control or individual YTHDF2 fragment to establish stable cell lines (Figure 5F). RT-qPCR analysis showing EBV ZTA and RTA mRNA levels in these cell lines upon lytic induction by anti-IgG treatment for 24 and 48 hrs. The value of vector control at 24 hrs was set as 1. (C) Akata (EBV+) cells were transduced with lentiviruses carrying vector control, Halo-tag, WT YTHDF2 or individual YTHDF2 fragment to establish stable cell lines. Western Blot analysis showing Halo, YTHDF2 fragments and EBV protein expression levels in these cell lines upon lytic induction by anti-IgG treatment for 24 and 48 hrs. Shorter and longer exposures were included to show the differences in protein levels. (D) Total mRNA was extracted from cells treated in panel (C). EBV ZTA and RTA mRNA levels were analyzed by RT-qPCR. The value of vector control at 24 hrs was set as 1 Results from three biological replicates are presented. Error bars indicate ±SD. N.S., not significant; ***, p< 0.001.

**Figure S10.**
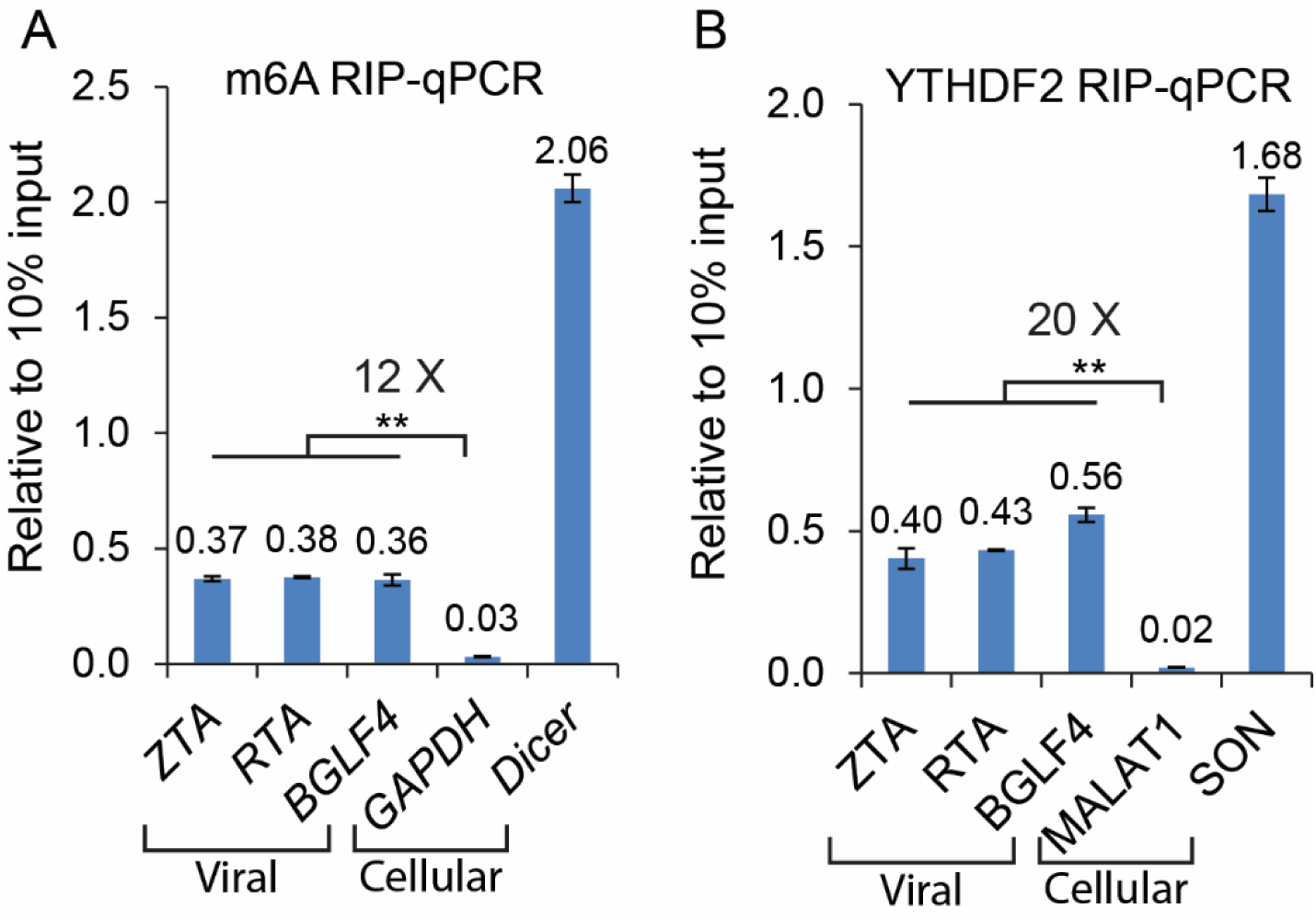
YTHDF2 binds to EBV transcripts and viral RNAs contain m^6^A modifications. See also Figure 6. Akata (EBV+) cells were lytically induced by IgG-cross linking for 24 hrs. (A) Total RNA was subjected to m^6^A RIP, followed by RT-qPCR using indicated primers. Values are displayed as fold change over 10% input. GAPDH and Dicer are cellular negative and positive controls, respectively. (B) Cell lysate was collected to detect YTHDF2 binding of viral RNAs by RIP-qPCR. Values are displayed as fold change over 10% input. MALAT1 and SON are cellular negative and positive controls, respectively. Results from three biological replicates are presented. Error bars indicate ±SD. **, p< 0.01.

**Figure S11.**
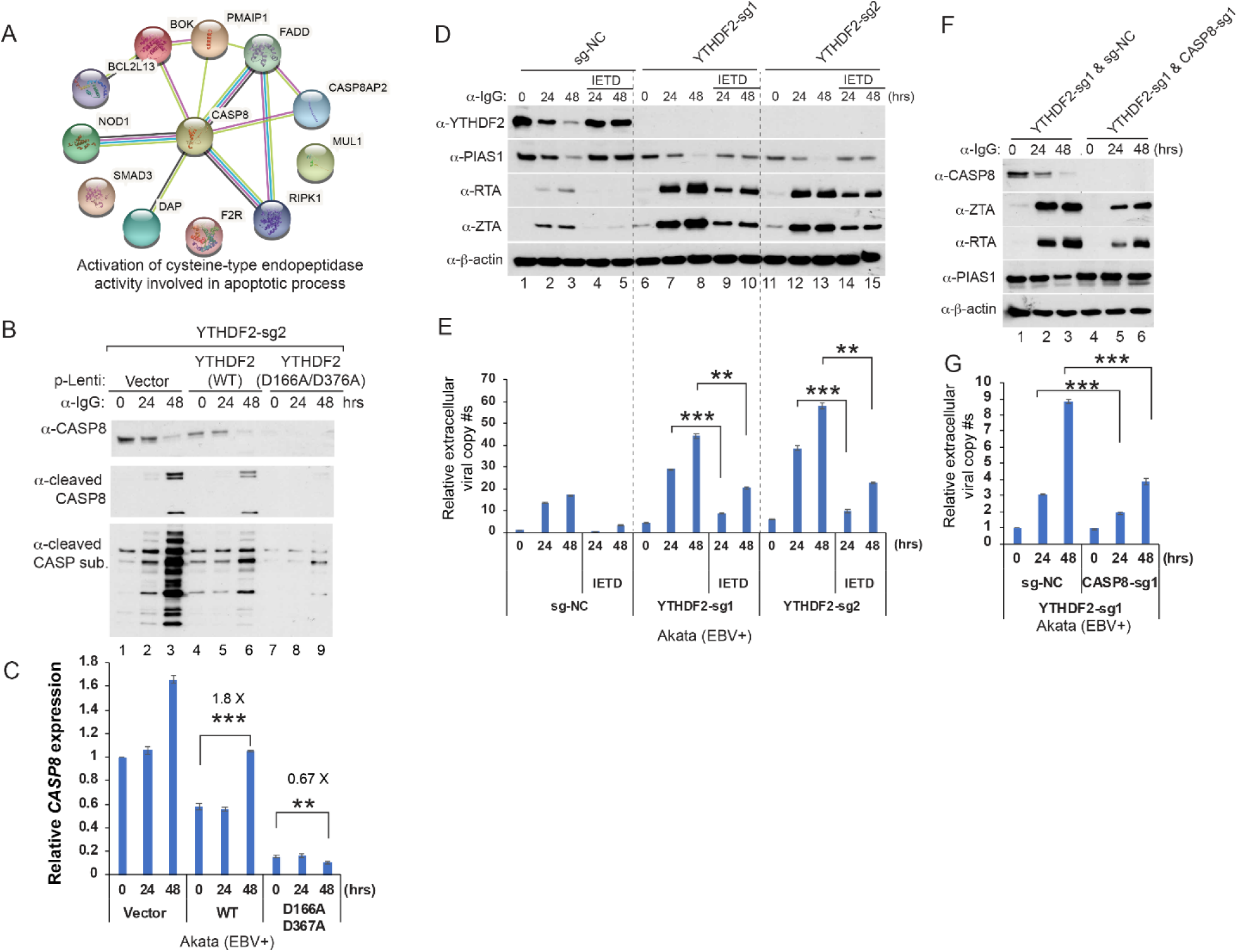
YTHDF2 cleavage or depletion promotes *CASP8* expression and caspase-8 inhibition limits EBV replication in YTHDF2-depleted cells. See also Figures 5-6. (A) A group of genes in the category of “*activation of cysteine-type endopeptidase activity involved in apoptotic process*” (also called “*caspase activation*”) were extracted from YTHDF2 target genes derived from YTHDF2 RIP-seq and PAR-CLIP-seq datasets (16, 51, 52) (B-C) YTHDF2 reconstitution suppresses caspase-8 expression and subsequent caspase activation. Akata (EBV+) YTHDF2-sg2 cells were reconstituted with WT or cleavage-resistant YTHDF2 (D166A/D367A) using lentiviral constructs. Western Blot analysis showing the levels for caspase-8 (CASP8), cleaved caspase-8, and cleaved caspase substrates (CASP sub.) in these cell lines upon IgG cross-linking as indicated (B). *CASP8* mRNA levels were analyzed by RT-qPCR using *CASP8* primers (C). The value of vector control at 0 hr was set as 1. (D-E) Caspase-8 inhibition suppress EBV replication in YTHDF2-depleted cells. Control and YTHDF2-depleted Akata (EBV+) cells were either untreated or pretreated with caspase-8 inhibitor (Z-IETD-FMK, 50 μM) for 1 hr and then anti-IgG antibody was added for 0 to 48 hrs as indicated. Western Blot showing the protein levels of EBV ZTA and RTA as indicated (D). Extracellular viral DNA was measured by qPCR using primers specific to BALF5 (E). The value of vector control at 0 hr was set as 1. (F-G) Caspase-8 depletion suppresses EBV replication in YTHDF2-depleted cells. YTHDF2-depleted Akata (EBV+) cells were transduced with lentivirus carrying control sgRNA or CASP8-sg1 to establish cell lines and then anti-IgG antibody was added for 0 to 48 hrs as indicated. Western Blot showing the protein levels of EBV ZTA and RTA as indicated (F). Extracellular viral DNA was measured by qPCR using primers specific to BALF5 (G). The value of vector control at 0 hr was set as 1. Results from three biological replicates are presented. Error bars indicate ±SD. **, p<0.01; ***, p<0.001.

**Figure S12.**
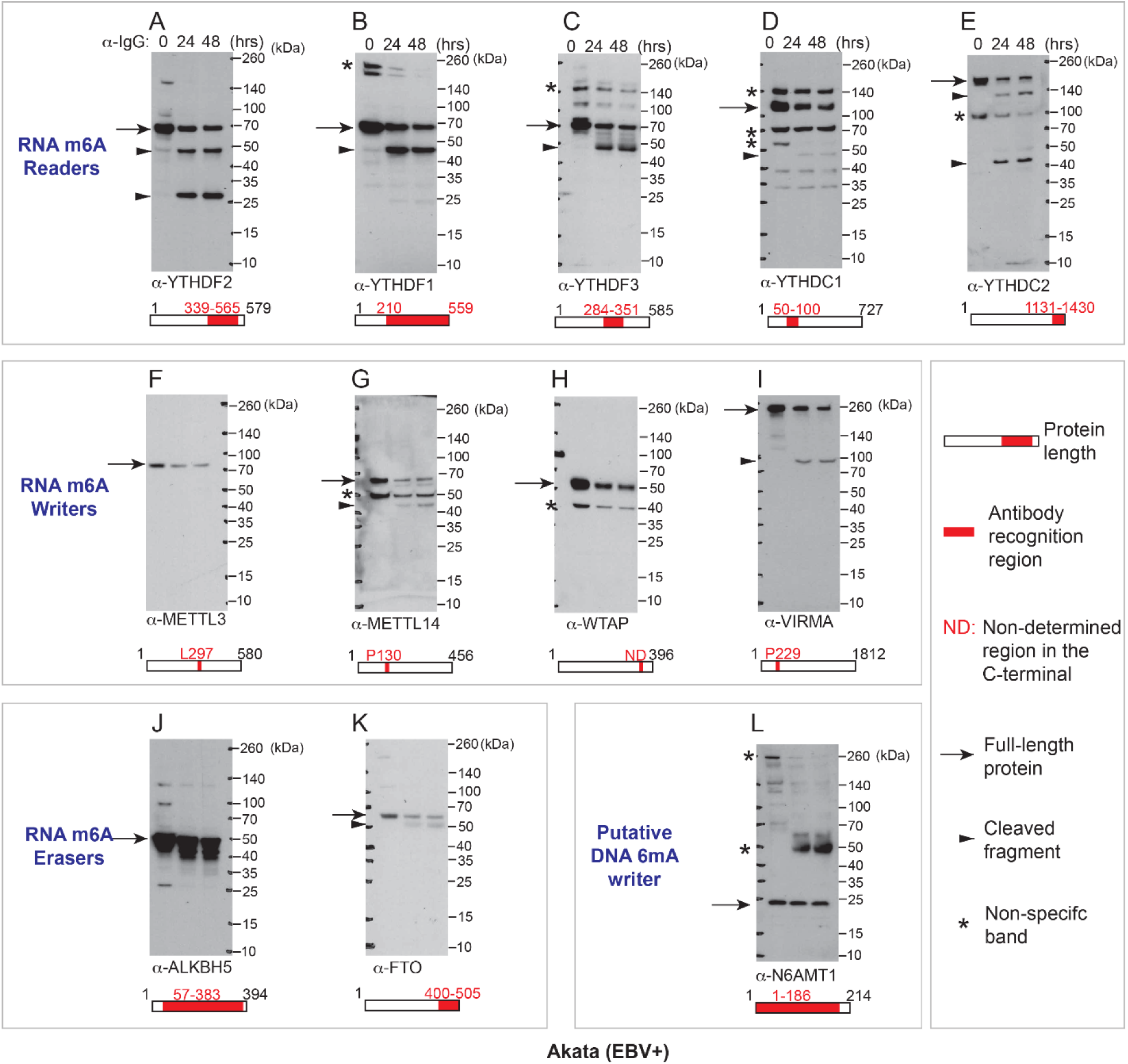
M^6^A readers, writers and erasers are cleaved upon lytic induction. See also Figure 7. Full blots for Figure 7B showing the generation of cleaved fragments for m^6^A readers, writers and erasers. Western Blot was performed using antibodies as indicated. N6AMT1 blot was included as a control.

**Figure S13.**
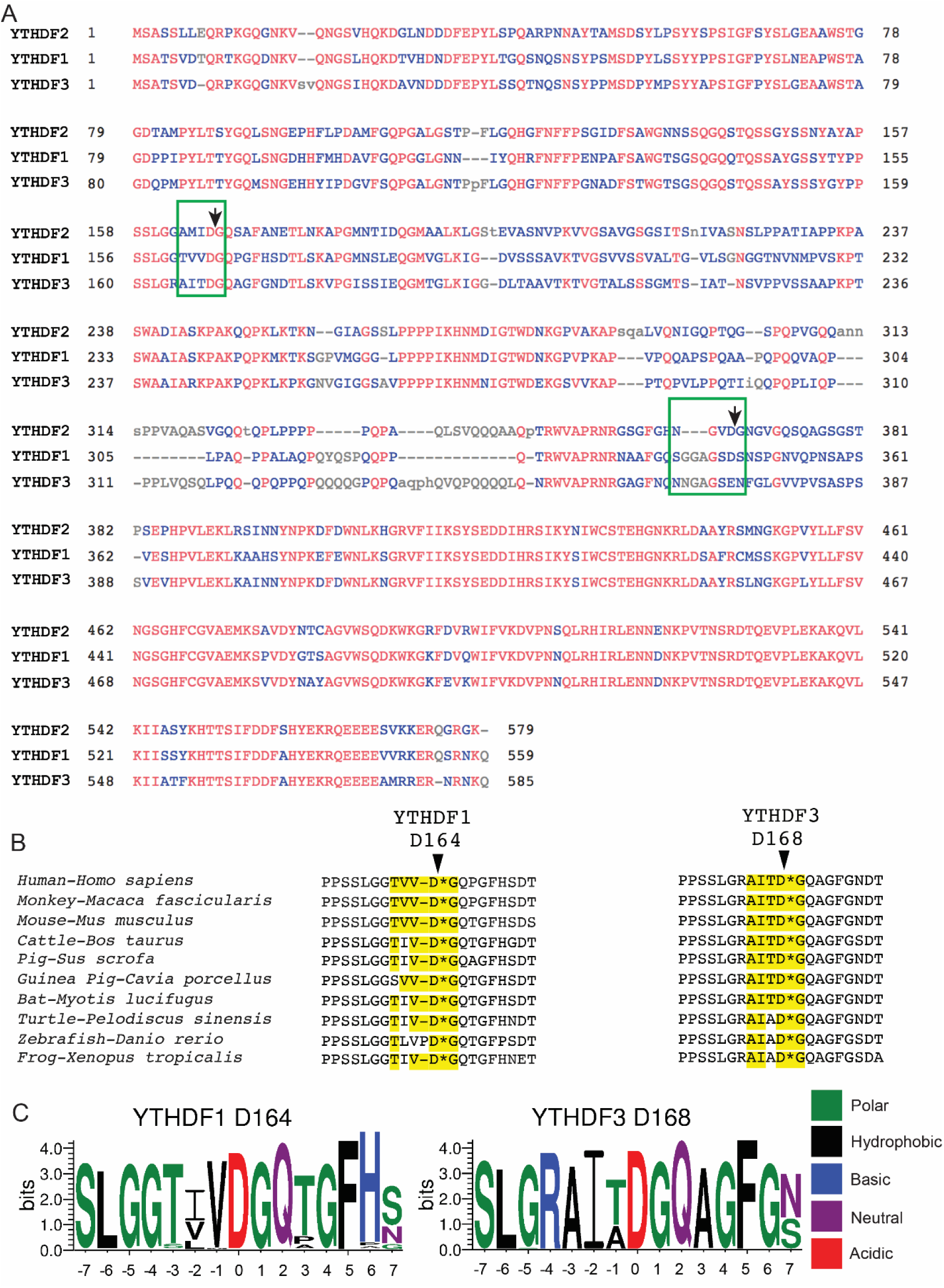
The conservation of YTHDF2, YTHDF1 and YTHDf3 cleavage sites. See also Figure 7 and Table S2. (A) Sequence alignment of YTHDF2 with YTHDF1 and YTHDF2 s using the Constraint-based Multiple Alignment Tool (COBALT). The corresponding cleavage motifs were highlighted by green boxes. (B) Sequence alignment of YTHDF2 sequences from 10 representative species using the Constraint-based Multiple Alignment Tool (COBALT). The cleavage motifs were highlighted by yellow color. (C) Motif analysis showing the conservation of YTHDF1-D164/YTHDF3-D168 and the surrounding amino acids. Amino acid sequences were extracted from 97 vertebrate species and motif logos were generated using WebLogo.

**Figure S14.**
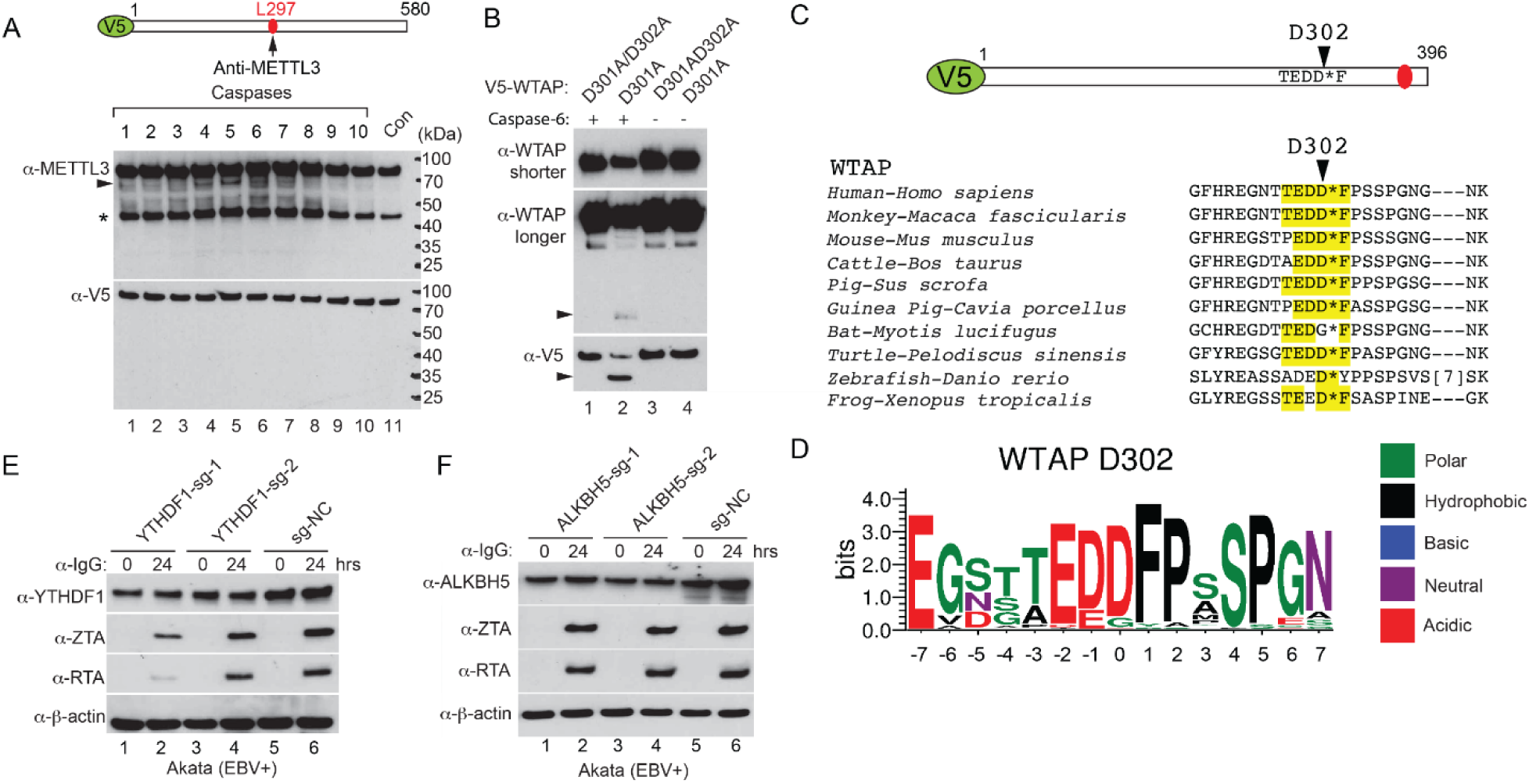
WTAP is cleaved on D302 and the depletion of YTHDF1 or ALKBH5 does not affect EBV protein expression. See also Figures 7, 8 and Table S2. (A) V5-METTL3 was incubated with individual caspase for 2 hrs at 37°C. Western Blot was performed using anti-METTL3 and anti-V5 antibodies as indicated. The locations of antibody recognition epitopes were labelled as indicated. The positions of weakly cleaved fragments were labelled by arrowhead. Star denotes non-specific bands. (B) V5-tagged WTAP D301A/D302A and D301A mutants were incubated with individual recombinant caspase for 2 hrs. Western Blot was performed using antibodies as indicated. Arrowheads denote cleaved fragments. (C) Sequence alignment of WTAP sequences from 10 representative species using the Constraint-based Multiple Alignment Tool (COBALT). The cleavage motifs were highlighted by yellow color. (D) Motif analysis showing the conservation of the WTAP D302 and the surrounding amino acids. Amino acid sequences were extracted from 97 vertebrate species and motif logos were generated using WebLogo. (E-F) Akata (EBV+) cells were used to establish stable cell lines using 2 different guide RNA constructs targeting YTHDF1 (D) and ALKBH5 (E) and a non-targeting control (sg-NC). The cells were untreated or lytically induced with anti-IgG-mediated BCR activation. Cellular and viral protein expression levels were monitored by Western Blot using antibodies as indicated.

**Figure S15.**
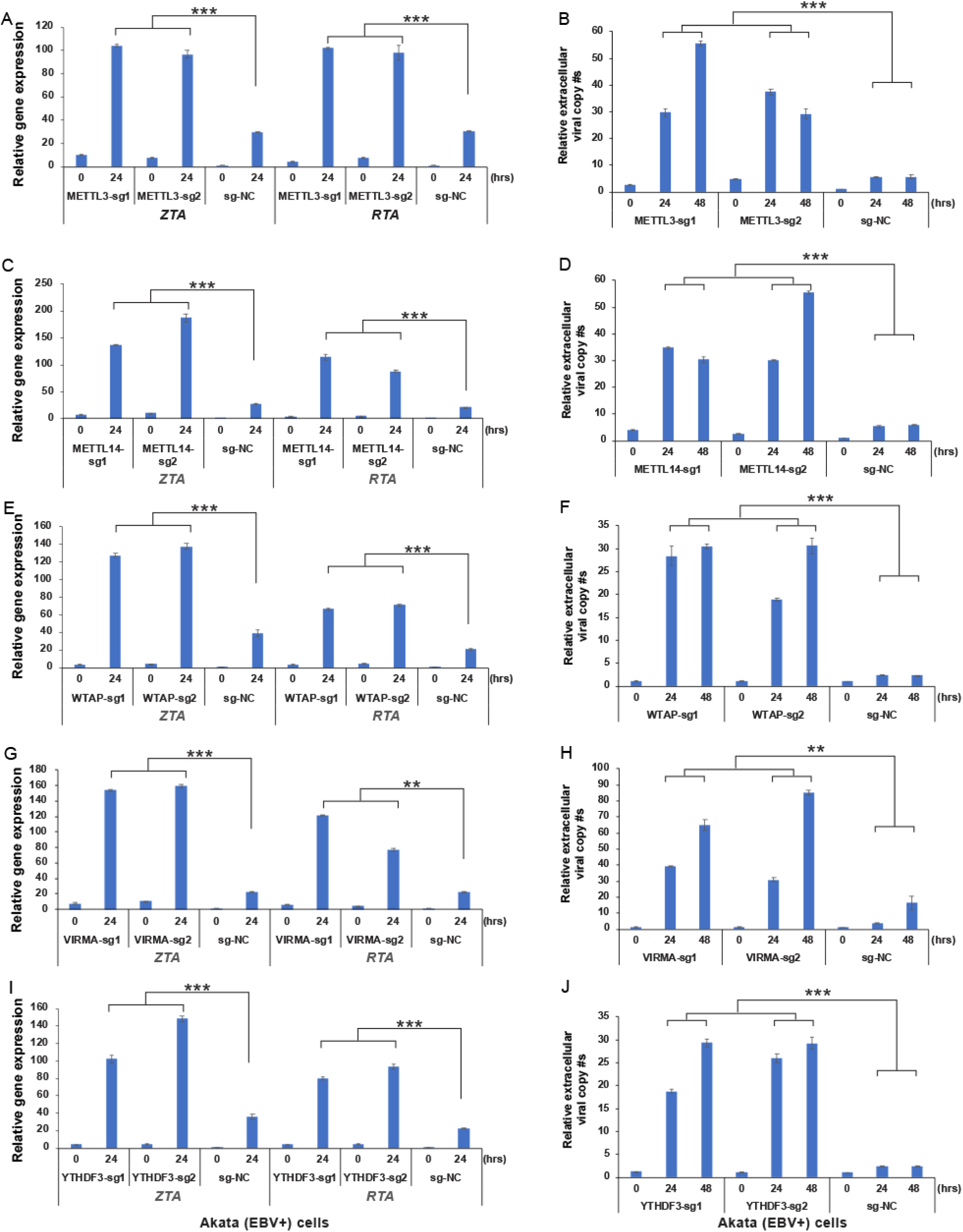
Depletion of m^6^A writers and reader YTHDF3 promotes EBV reactivation. See also Figure 8. Akata (EBV+) cells were used to establish stable cell lines using 2-3 different guide RNA constructs targeting METTL3 (A and B), METTL14 (C and D), WTAP (E and F), VIRMA (G and H) and YTHDF3 (I and J) and a non-targeting control (sg-NC). The cells were untreated or lytically induced with anti-IgG-mediated BCR activation for 24 or 48 hrs. EBV *ZTA* and *RTA* mRNA expression levels were monitored by RT-qPCR (A, C, E, G and I). Extracellular viral DNA was measured by qPCR using primers specific to BALF5 (B, D, F, H and J). The value of vector control at 0 hr was set as 1. Results from three biological replicates are presented. Error bars indicate ±SD. **, p<0.01; ***, p<0.001.

**Figure S16.**
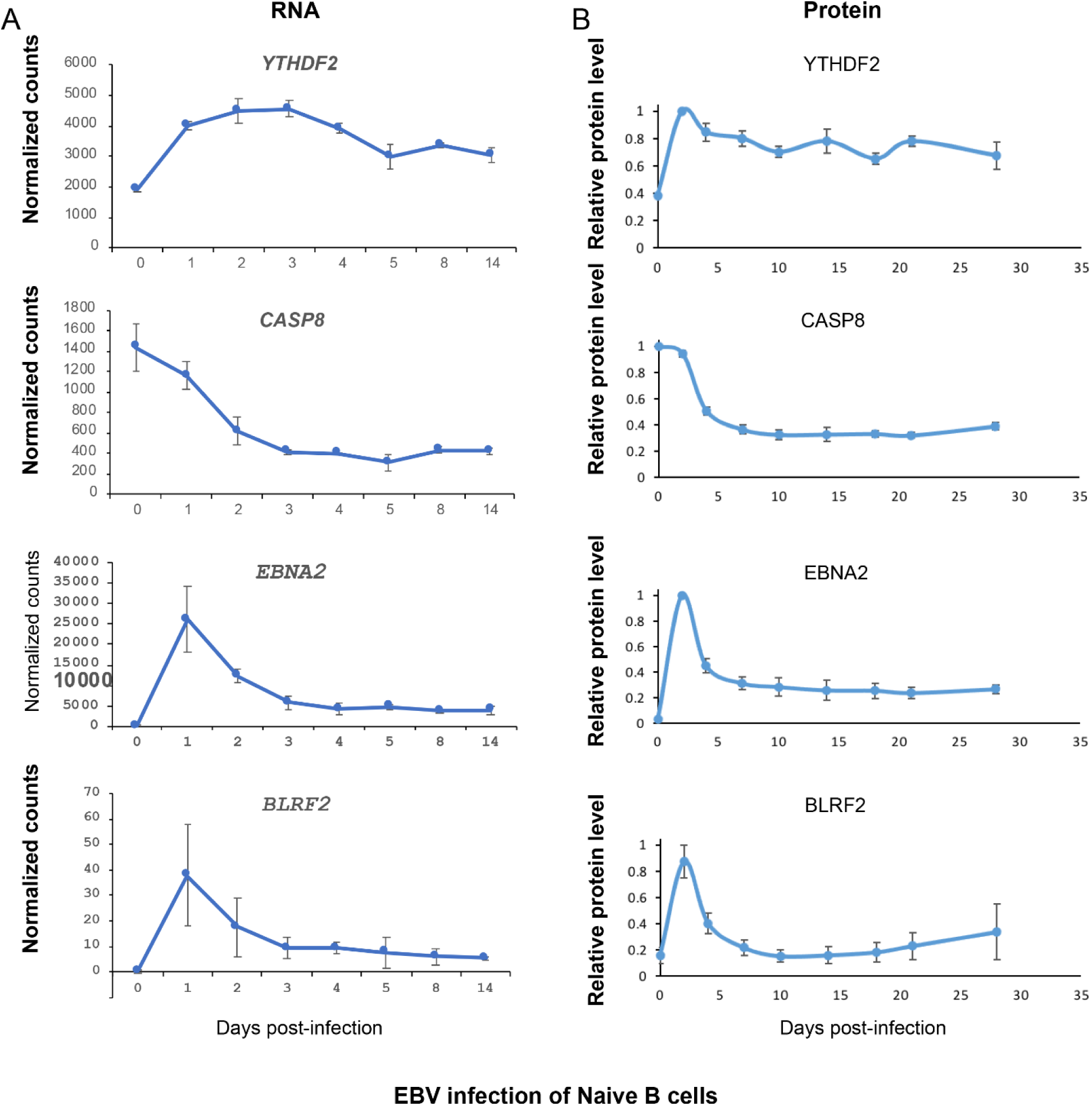
The expression of YTHDF2 negatively correlates with the mRNA and protein expression of CASP8, EBV EBNA2 and BLRF2 upon EBV infection of primary human B cells. The RNA (A) and protein (B) levels of YTHDF2, CASP8, EBV EBNA2 and BRLF2 were analyzed for EBV infection of primary human B cells. The data were extracted from the transcriptomic (http://ebv-b.helmholtz-muenchen.de/) (57) and proteomic analyses (56), respectively.

Table S1. Primers, templates and key resources used in this study.

Table S2. Table S2. Caspase cleavage motif sequences (corresponding to cleavage sites in human YTHDF2, YTHDF1, YTHDF3 and WTAP) extracted from vertebrate species. Related to Figures 4, S13 and S14

Table S3. M^6^A motif sequences (corresponding to putative m^6^A motifs in human *CASP8*-Exon 7) extracted from vertebrate species. Related to Figure 6.

## References

1. Young LS, Rickinson AB. Epstein-Barr virus: 40 years on. Nat Rev Cancer. 2004;4(10):757–68.

2. Young LS, Yap LF, Murray PG. Epstein-Barr virus: more than 50 years old and still providing surprises. Nat Rev Cancer. 2016;16(12):789–802.

3. Cohen JI, Fauci AS, Varmus H, Nabel GJ. Epstein-Barr virus: an important vaccine target for cancer prevention. Sci Transl Med. 2011;3(107):107fs7.

4. De Leo A, Calderon A, Lieberman PM. Control of Viral Latency by Episome Maintenance Proteins. Trends Microbiol. 2020;28(2):150–62.

5. Kenney SC, Mertz JE. Regulation of the latent-lytic switch in Epstein-Barr virus. Semin Cancer Biol. 2014;26:60–8.

6. Lv DW, Zhang K, Li R. Interferon regulatory factor 8 regulates caspase-1 expression to facilitate Epstein-Barr virus reactivation in response to B cell receptor stimulation and chemical induction. PLoS pathogens. 2018;14(1):e1006868.

7. Zhang K, Lv DW, Li R. B Cell Receptor Activation and Chemical Induction Trigger Caspase-Mediated Cleavage of PIAS1 to Facilitate Epstein-Barr Virus Reactivation. Cell reports. 2017;21(12):3445–57.

8. Lieberman PM. Keeping it quiet: chromatin control of gammaherpesvirus latency. Nat Rev Microbiol. 2013;11(12):863–75.

9. Fu DX, Tanhehco Y, Chen J, Foss CA, Fox JJ, Chong JM, et al. Bortezomib-induced enzyme-targeted radiation therapy in herpesvirus-associated tumors. Nature medicine. 2008;14(10):1118–22.

10. Meyer KD, Jaffrey SR. Rethinking m(6)A Readers, Writers, and Erasers. Annual review of cell and developmental biology. 2017;33:319–42.

11. Manners O, Baquero-Perez B, Whitehouse A. m(6)A: Widespread regulatory control in virus replication. Biochimica et biophysica acta Gene regulatory mechanisms. 2018.

12. Williams GD, Gokhale NS, Horner SM. Regulation of Viral Infection by the RNA Modification N6-methyladenosine. Annual review of virology. 2019;29(6(1)):235–53.

13. Liu J, Yue Y, Han D, Wang X, Fu Y, Zhang L, et al. A METTL3-METTL14 complex mediates mammalian nuclear RNA N6-adenosine methylation. Nature chemical biology. 2014;10(2):93–5.

14. Zheng G, Dahl JA, Niu Y, Fedorcsak P, Huang CM, Li CJ, et al. ALKBH5 is a mammalian RNA demethylase that impacts RNA metabolism and mouse fertility. Molecular cell. 2013;49(1):18–29.

15. Jia G, Fu Y, Zhao X, Dai Q, Zheng G, Yang Y, et al. N6-methyladenosine in nuclear RNA is a major substrate of the obesity-associated FTO. Nature chemical biology. 2011;7(12):885–7.

16. Wang X, Lu Z, Gomez A, Hon GC, Yue Y, Han D, et al. N6-methyladenosine-dependent regulation of messenger RNA stability. Nature. 2014;505(7481):117–20.

17. Shi H, Wang X, Lu Z, Zhao BS, Ma H, Hsu PJ, et al. YTHDF3 facilitates translation and decay of N(6)-methyladenosine-modified RNA. Cell research. 2017;27(3):315–28.

18. Du H, Zhao Y, He J, Zhang Y, Xi H, Liu M, et al. YTHDF2 destabilizes m(6)A-containing RNA through direct recruitment of the CCR4-NOT deadenylase complex. Nature communications. 2016;7:12626.

19. Lasman L, Krupalnik V, Viukov S, Mor N, Aguilera-Castrejon A, Schneir D, et al. Context-dependent functional compensation between Ythdf m(6)A reader proteins. Genes & development. 2020.

20. Zaccara S, Jaffrey SR. A Unified Model for the Function of YTHDF Proteins in Regulating m(6)A-Modified mRNA. Cell. 2020;181(7):1582–95 e18.

21. Fu Y, Zhuang X. m(6)A-binding YTHDF proteins promote stress granule formation. Nature chemical biology. 2020;16(9):955–63.

22. Ries RJ, Zaccara S, Klein P, Olarerin-George A, Namkoong S, Pickering BF, et al. m(6)A enhances the phase separation potential of mRNA. Nature. 2019;571(7765):424–8.

23. Wang J, Wang L, Diao J, Shi YG, Shi Y, Ma H, et al. Binding to m(6)A RNA promotes YTHDF2-mediated phase separation. Protein Cell. 2020;11(4):304–7.

24. Xia TL, Li X, Wang X, Zhu YJ, Zhang H, Cheng W, et al. N(6)-methyladenosine-binding protein YTHDF1 suppresses EBV replication and promotes EBV RNA decay. EMBO Rep. 2021:e50128.

25. Lang F, Singh RK, Pei Y, Zhang S, Sun K, Robertson ES. EBV epitranscriptome reprogramming by METTL14 is critical for viral-associated tumorigenesis. PLoS pathogens. 2019;15(6):e1007796.

26. Dai DL, Li X, Wang L, Xie C, Jin Y, Zeng MS, et al. Identification of an N6-methyladenosine-mediated positive feedback loop that promotes Epstein-Barr virus infection. J Biol Chem. 2021:100547.

27. Zheng X, Wang J, Zhang X, Fu Y, Peng Q, Lu J, et al. RNA m(6) A methylation regulates virus-host interaction and EBNA2 expression during Epstein-Barr virus infection. Immun Inflamm Dis. 2021.

28. Tan B, Liu H, Zhang S, da Silva SR, Zhang L, Meng J, et al. Viral and cellular N(6)-methyladenosine and N(6),2’-O-dimethyladenosine epitranscriptomes in the KSHV life cycle. Nature microbiology. 2018;3(1):108–20.

29. Hesser CR, Karijolich J, Dominissini D, He C, Glaunsinger BA. N6-methyladenosine modification and the YTHDF2 reader protein play cell type specific roles in lytic viral gene expression during Kaposi’s sarcoma-associated herpesvirus infection. PLoS pathogens. 2018;14(4):e1006995.

30. Ye F, Chen ER, Nilsen TW. Kaposi’s Sarcoma-Associated Herpesvirus Utilizes and Manipulates RNA N(6)-Adenosine Methylation To Promote Lytic Replication. Journal of virology. 2017;91(16).

31. Baquero-Perez B, Antanaviciute A, Yonchev ID, Carr IM, Wilson SA, Whitehouse A. The Tudor SND1 protein is an m(6)A RNA reader essential for replication of Kaposi’s sarcoma-associated herpesvirus. eLife. 2019;8.

32. Rubio RM, Depledge DP, Bianco C, Thompson L, Mohr I. RNA m(6) A modification enzymes shape innate responses to DNA by regulating interferon beta. Genes & development. 2018;32(23-24):1472–84.

33. De Leo A, Chen HS, Hu CA, Lieberman PM. Deregulation of KSHV latency conformation by ER-stress and caspase-dependent RAD21-cleavage. PLoS Pathog. 2017;13(8):e1006596.

34. Tabtieng T, Degterev A, Gaglia MM. Caspase-Dependent Suppression of Type I Interferon Signaling Promotes Kaposi’s Sarcoma-Associated Herpesvirus Lytic Replication. Journal of virology. 2018;92(10).

35. Burton EM, Goldbach-Mansky R, Bhaduri-McIntosh S. A promiscuous inflammasome sparks replication of a common tumor virus. Proceedings of the National Academy of Sciences of the United States of America. 2020;117(3):1722–30.

36. Julien O, Wells JA. Caspases and their substrates. Cell Death Differ. 2017;24(8):1380–9.

37. Lv DW, Zhong J, Zhang K, Pandey A, Li R. Understanding Epstein-Barr Virus Life Cycle with Proteomics: A Temporal Analysis of Ubiquitination During Virus Reactivation. OMICS: A Journal of Integrative Biology. 2017;21(1):27–37.

38. Zhang K, Lv DW, Li R. Conserved Herpesvirus Protein Kinases Target SAMHD1 to Facilitate Virus Replication. Cell reports. 2019;28(2):449–59 e5.

39. Kumar S, van Raam BJ, Salvesen GS, Cieplak P. Caspase cleavage sites in the human proteome: CaspDB, a database of predicted substrates. PloS one. 2014;9(10):e110539.

40. Yang J, Zhang Y. I-TASSER server: new development for protein structure and function predictions. Nucleic acids research. 2015;43(W1):W174–81.

41. Zhang C, Freddolino PL, Zhang Y. COFACTOR: improved protein function prediction by combining structure, sequence and protein-protein interaction information. Nucleic acids research. 2017;45(W1):W291–W9.

42. Li F, Zhao D, Wu J, Shi Y. Structure of the YTH domain of human YTHDF2 in complex with an m(6)A mononucleotide reveals an aromatic cage for m(6)A recognition. Cell research. 2014;24(12):1490–2.

43. Zhu T, Roundtree IA, Wang P, Wang X, Wang L, Sun C, et al. Crystal structure of the YTH domain of YTHDF2 reveals mechanism for recognition of N6-methyladenosine. Cell research. 2014;24(12):1493–6.

44. Pettersen EF, Goddard TD, Huang CC, Couch GS, Greenblatt DM, Meng EC, et al. UCSF Chimera--a visualization system for exploratory research and analysis. J Comput Chem. 2004;25(13):1605–12.

45. Feng Y, Daley-Bauer LP, Roback L, Guo H, Koehler HS, Potempa M, et al. Caspase-8 restricts antiviral CD8 T cell hyperaccumulation. Proceedings of the National Academy of Sciences of the United States of America. 2019;116(30):15170–7.

46. Kaiser WJ, Upton JW, Long AB, Livingston-Rosanoff D, Daley-Bauer LP, Hakem R, et al. RIP3 mediates the embryonic lethality of caspase-8-deficient mice. Nature. 2011;471(7338):368–72.

47. Tan B, Gao SJ. The RNA Epitranscriptome of DNA Viruses. Journal of virology. 2018;92(22).

48. Fei Q, Zou Z, Roundtree IA, Sun HL, He C. YTHDF2 promotes mitotic entry and is regulated by cell cycle mediators. PLoS Biol. 2020;18(4):e3000664.

49. Dominissini D, Moshitch-Moshkovitz S, Schwartz S, Salmon-Divon M, Ungar L, Osenberg S, et al. Topology of the human and mouse m6A RNA methylomes revealed by m6A-seq. Nature. 2012;485(7397):201–6.

50. Meyer KD, Saletore Y, Zumbo P, Elemento O, Mason CE, Jaffrey SR. Comprehensive analysis of mRNA methylation reveals enrichment in 3’ UTRs and near stop codons. Cell. 2012;149(7):1635–46.

51. Liu H, Flores MA, Meng J, Zhang L, Zhao X, Rao MK, et al. MeT-DB: a database of transcriptome methylation in mammalian cells. Nucleic acids research. 2015;43(Database issue):D197–203.

52. Liu H, Wang H, Wei Z, Zhang S, Hua G, Zhang SW, et al. MeT-DB V2.0: elucidating context-specific functions of N6-methyl-adenosine methyltranscriptome. Nucleic acids research. 2018;46(D1):D281–D7.

53. Linder B, Grozhik AV, Olarerin-George AO, Meydan C, Mason CE, Jaffrey SR. Single-nucleotide-resolution mapping of m6A and m6Am throughout the transcriptome. Nature methods. 2015;12(8):767–72.

54. Gokhale NS, McIntyre ABR, Mattocks MD, Holley CL, Lazear HM, Mason CE, et al. Altered m(6)A Modification of Specific Cellular Transcripts Affects Flaviviridae Infection. Molecular cell. 2020;77(3):542–55 e8.

55. Mahrus S, Trinidad JC, Barkan DT, Sali A, Burlingame AL, Wells JA. Global sequencing of proteolytic cleavage sites in apoptosis by specific labeling of protein N termini. Cell. 2008;134(5):866–76.

56. Wang LW, Shen H, Nobre L, Ersing I, Paulo JA, Trudeau S, et al. Epstein-Barr-Virus-Induced One-Carbon Metabolism Drives B Cell Transformation. Cell Metab. 2019;30(3):539–55 e11.

57. Mrozek-Gorska P, Buschle A, Pich D, Schwarzmayr T, Fechtner R, Scialdone A, et al. Epstein-Barr virus reprograms human B lymphocytes immediately in the prelatent phase of infection. Proceedings of the National Academy of Sciences of the United States of America. 2019;116(32):16046–55.

58. Wei J, He C. Chromatin and transcriptional regulation by reversible RNA methylation. Curr Opin Cell Biol. 2021;70:109–15.

59. Wang Y, Li Y, Yue M, Wang J, Kumar S, Wechsler-Reya RJ, et al. N(6)-methyladenosine RNA modification regulates embryonic neural stem cell self-renewal through histone modifications. Nat Neurosci. 2018;21(2):195–206.

60. Chen J, Zhang YC, Huang C, Shen H, Sun B, Cheng X, et al. m(6)A Regulates Neurogenesis and Neuronal Development by Modulating Histone Methyltransferase Ezh2. Genomics Proteomics Bioinformatics. 2019;17(2):154–68.

61. Toth Z, Maglinte DT, Lee SH, Lee HR, Wong LY, Brulois KF, et al. Epigenetic analysis of KSHV latent and lytic genomes. PLoS pathogens. 2010;6(7):e1001013.

62. Murata T, Kondo Y, Sugimoto A, Kawashima D, Saito S, Isomura H, et al. Epigenetic histone modification of Epstein-Barr virus BZLF1 promoter during latency and reactivation in Raji cells. Journal of virology. 2012;86(9):4752–61.

63. Zerby D, Chen CJ, Poon E, Lee D, Shiekhattar R, Lieberman PM. The amino-terminal C/H1 domain of CREB binding protein mediates zta transcriptional activation of latent Epstein-Barr virus. Mol Cell Biol. 1999;19(3):1617–26.

64. Liu J, Dou X, Chen C, Chen C, Liu C, Xu MM, et al. N (6)-methyladenosine of chromosome-associated regulatory RNA regulates chromatin state and transcription. Science. 2020;367(6477):580–6.

65. Xu W, Li J, He C, Wen J, Ma H, Rong B, et al. METTL3 regulates heterochromatin in mouse embryonic stem cells. Nature. 2021;591(7849):317–21.

66. Sanjana NE, Shalem O, Zhang F. Improved vectors and genome-wide libraries for CRISPR screening. Nat Methods. 2014;11(8):783–4.

67. Golden RJ, Chen B, Li T, Braun J, Manjunath H, Chen X, et al. An Argonaute phosphorylation cycle promotes microRNA-mediated silencing. Nature. 2017;542(7640):197–202.

68. Li R, Liao G, Nirujogi RS, Pinto SM, Shaw PG, Huang TC, et al. Phosphoproteomic Profiling Reveals Epstein-Barr Virus Protein Kinase Integration of DNA Damage Response and Mitotic Signaling. PLoS pathogens. 2015;11(12):e1005346.

69. Hubisz MJ, Pollard KS, Siepel A. PHAST and RPHAST: phylogenetic analysis with space/time models. Brief Bioinform. 2011;12(1):41–51.

70. Crooks GE, Hon G, Chandonia JM, Brenner SE. WebLogo: a sequence logo generator. Genome Res. 2004;14(6):1188–90.

71. Wagih O. ggseqlogo: a versatile R package for drawing sequence logos. Bioinformatics. 2017;33(22):3645–7.

